# Lineage tracing on transcriptional landscapes links state to fate during differentiation

**DOI:** 10.1101/467886

**Authors:** Caleb Weinreb, Alejo Rodriguez-Fraticelli, Fernando Camargo, Allon M Klein

## Abstract

A challenge in stem cell biology is to associate molecular differences among progenitor cells with their capacity to generate mature cell types. Though the development of single cell assays allows for the capture of progenitor cell states in great detail, these assays cannot definitively link cell states to their long-term fate. Here, we use expressed DNA barcodes to clonally trace single cell transcriptomes dynamically during differentiation and apply this approach to the study of hematopoiesis. Our analysis identifies functional boundaries of cell potential early in the hematopoietic hierarchy and locates them on a continuous transcriptional landscape. We reconstruct a developmental hierarchy showing separate ontogenies for granulocytic subtypes and two routes to monocyte differentiation that leave a persistent imprint on mature cells. Finally, we use our approach to benchmark methods of dynamic inference from single-cell snapshots, and provide evidence of strong early fate biases dependent on cellular properties hidden from single-cell RNA sequencing.

## Introduction

To generate and maintain multicellular organisms, cells must differentiate into distinct types with varied functions and molecular components. Progenitor cells progress through a hierarchy of fate decisions, refining their identity until reaching a functional end state. The gold standard for inferring the relationship between progenitors and their offspring is fate mapping, where a subset of progenitors is labeled, typically using genetic approaches that mark cells expressing defined marker genes. The fate of the labeled cells and their progeny is profiled at a later time point^1^. Fate maps are key to understanding and even controlling differentiation across a broad range of systems^2^.

Recently, whole-genome approaches for profiling cells by single cell RNA sequencing (scSeq) opened up a complementary approach to understand developmental relationships. scSeq can capture mature cell types alongside all stages of cell differentiation, revealing a ‘state map’ in gene expression space. Specialized algorithms applied to these state maps offer hypotheses for the hierarchy of cell states^3^ and their gene expression dynamics over time^4-7^. Unlike fate mapping approaches, scSeq state mapping can be carried out without prior genetic manipulation, and without being limited by the specificity of transgene expression within the progenitor cell pool^2^.

Neither state or fate mapping alone, however, provide a complete view of differentiation processes. Whereas scSeq offers a far higher resolution of cell states than fate mapping, it cannot not definitively link the detailed states of progenitors to their ultimate fate, because cells are destroyed in the process of measurement. scSeq data does not directly report the stages at which progenitor cells become committed to one or more fates or how many distinct paths might lead cells to the same end states. In addition, the sparse and high-dimensional nature of scSeq means that there may be more than one approach to constructing cell state trajectories from the same data^4^. There is a need for approaches that can link the detailed state of a cell to its long-term dynamic behavior and ultimate fate.

In this paper we integrate measurements of cell lineage with scSeq, using the mouse hematopoietic system as a model of fate choice. In adults, hematopoietic progenitor cells (HPCs) reside in the bone marrow and maintain steady-state blood production. Transplantation studies over several decades have led to the prevailing model of hematopoiesis as a branching hierarchy with defined oligo-potent intermediates^8^. But recent state maps from scSeq^9^, as well as clonal studies using barcodes^10^ and single cell culture^11^, suggest that the traditional intermediate cell types are internally heterogeneous in state and fate potential, with HPCs lying along a continuum of states rather than a well-defined hierarchy. Reconciling these views requires tracking the dynamics of individual lineages on the continuous landscape of HPC states defined by scSeq^12^. We explore an experimental design for capturing the state of a cell at the whole-transcriptome level, and its clonal fate at a later time point, simultaneously across thousands of cells in different states.

## RESULTS

### A simultaneous assay of clonal states and fates

Our strategy for simultaneously capturing transcriptional cell state and fate is to genetically barcode a population of progenitor cells, allow them to divide briefly, sample some immediately for scSeq profiling, and sample the remainder at later time points^13^. This approach will provide data for three types of clonal relationships (Fig. 1a): (1) measurements at the earliest time points will potentially capture two sister cells (mothers) after 1 or 2 rounds of cell division, revealing their initial transcriptional state and relatedness; (2) additional measurements at later time points, e.g. after differentiation, will identify the lineage fate of daughter(s) and allow comparison with the state of the clonally-related early mother, and (3) measurements of multiple differentiated daughters will reveal the clonal coupling between different lineages. If recently-divided mother cells (type 1) are transcriptionally similar, then pairs of clonally-related mother-daughters sampled across time (type 2) should reveal how single cell gene expression changes over time and with differentiation. Unlike conventional lineage tracing^2^, this approach can map the fate of cells from a continuous landscape of starting states. Further, it does not require a labeling approach targeted at specific progenitor cells (cf. ^14^).

**Figure 1:**
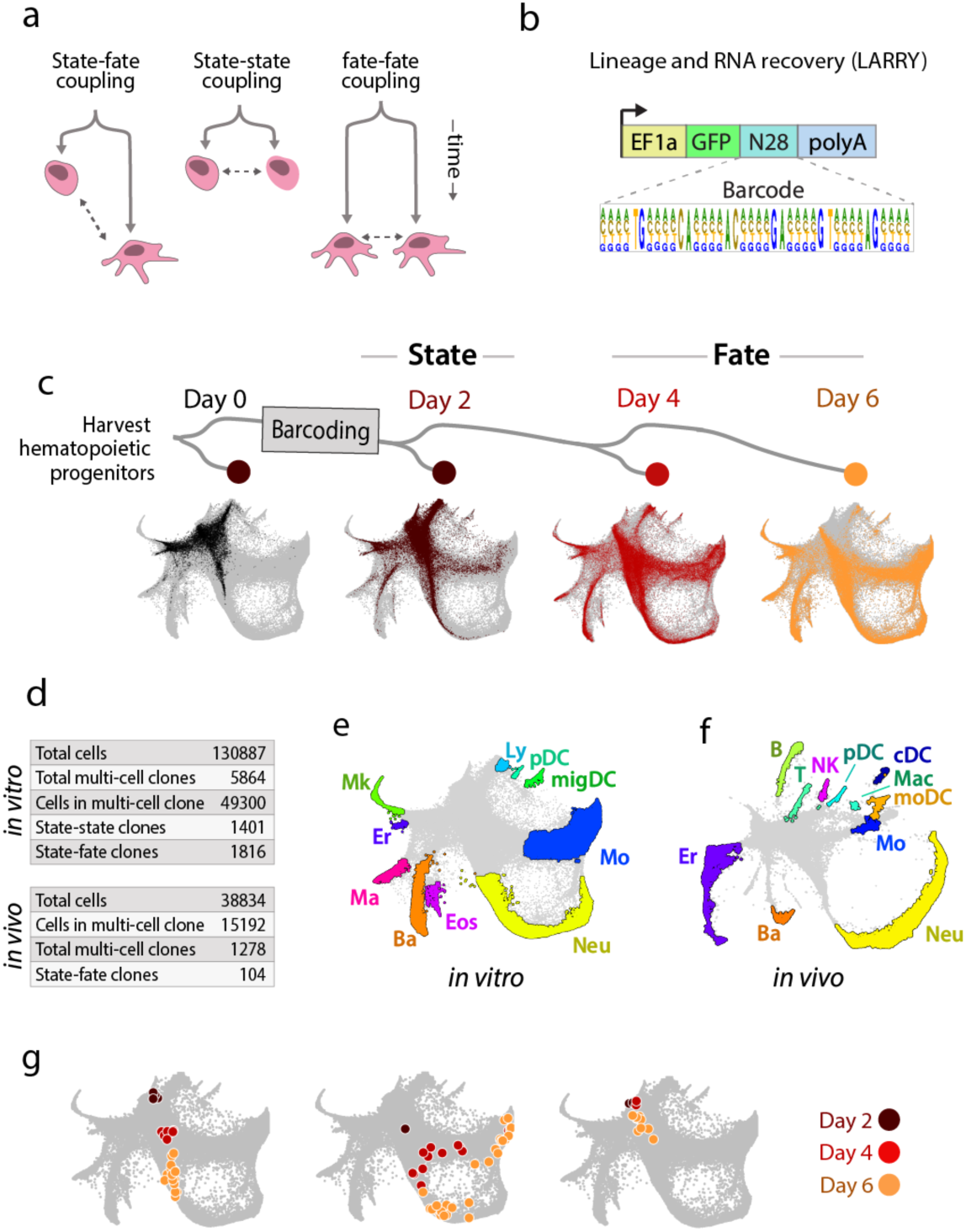
Tracking thousands of lineages across hematopoietic differentiation in vitro. (a) Experimental designs for tracking differentiation dynamics by analysis of sister cells. “State” refers to the gene expression state of a cell before differentiation has occurred. “Fate” corresponds to a cell’s lineage fate after differentiation. (b) Lentiviral construct for delivering a unique and heritable barcode that is expressed in the 3’ UTR of GFP and detectable using scSeq. (c) Tracking clones of hematopoietic progenitors over multiple time points in primary culture. Plots indicate the states of single-cell transcriptomes from each time point using a force-layout (SPRING) algorithm. (d) Total numbers of cells and clones sampled. (e-f) Annotation of cell types that appeared after 6 days of culture (e) or one week in vivo (f), based on gene expression [Ly=lymphoid precursor, Mk=megakaryocyte, Er=erythrocyte, Ma=mast cell, Ba=basophil, Eos=eosinophil, Neu=neutrophil, Mo=monocyte, Mac=macrophage, DC=dendritic cell, migDC=migratory (ccr7+) DC, pDC=plasmacytoid DC, moDC=monocyte-derived DC, cDC=classical DC, NK=natural killer cells, T=T-cell, B=B-cell]. (g) Examples of clonal dynamics on the single cell landscape. Each plot shows a separate clone, with cells colored by time point and overlaid on the full dataset in gray. The clones demonstrate uni-lineage differentiation (left), multi-lineage differentiation (middle) and self-renewal (right).

We modified a classical strategy for clonal labeling by lentiviral delivery of inherited DNA barcodes^15,16^, to allow barcode detection using scSeq^17^. The barcode consists of a random 28-mer in the 3’ UTR of an enhanced green fluorescent protein (*eGFP*) transgene under control of a ubiquitous EF1*α* promoter (Fig 1b). Transcripts of eGFP are captured during scSeq, and the barcode is revealed through analysis of sequencing reads. The theoretical number of barcodes for this construct is 4^28^ (or ∼10^16^), though bottlenecks in molecular cloning limited the diversity to ∼10^6^ barcodes, sufficient to label ∼10^4^ cells in an experiment with <0.5% barcode overlap between clones. We refer to the barcoding construct as LARRY (Lineage And RNA RecoverY).

We tested LARRY on mouse embryonic stem (ES) cells and primary HPCs. After profiling by scSeq, one or more barcodes could be robustly detected in 93% of GFP+ cells (Supp Fig 1a-c). Specific barcode sequences overlapped rarely between replicate transduction experiments, at a frequency expected by chance for the library size (0.3% of 5000 barcodes appeared more than once). Virally transduced HPCs underwent transient aggregation but showed little difference in gene expression or lineage distribution compared to controls when analyzed by scSeq (Supp Fig 1d,e), consistent with previous reports^18^. Therefore, the approach provides an efficient method for simultaneously barcoding large numbers of cells for combined fate and state mapping.

To analyze HPC fate potential, we isolated a broad class of oligo-potent (Lin-Kit+, or LK) and multipotent progenitors (Lin-Sca1+Kit+ or LSK) cells (Supp Fig 2a,b) and plated them in media chosen to support broad multi-lineage differentiation (see Methods). Following transduction with LARRY, cells were cultured for two days to allow time for lentiviral integration and subsequent cell division. During this time the cells divided three times on average, with some division expected to occur before lentiviral integration. We then sampled half the cells (defining the ‘early state’) and profiled them by scSeq. The other half was re-plated and then sampled after two days (30% of cells) and four days (remaining cells) (Fig 1c). To benchmark differentiation in culture, we similarly transplanted barcoded progenitor cells into an irradiated host mouse to differentiate 1 week in vivo. Together, the samples included 130,887 cultured single cells, of which 38% belonged to a clone of two or more cells; as well as 38,834 cells before or after *in vivo* engraftment, with a similar fraction (39%) in multi-cell clones (Fig 1d).

**Figure 2:**
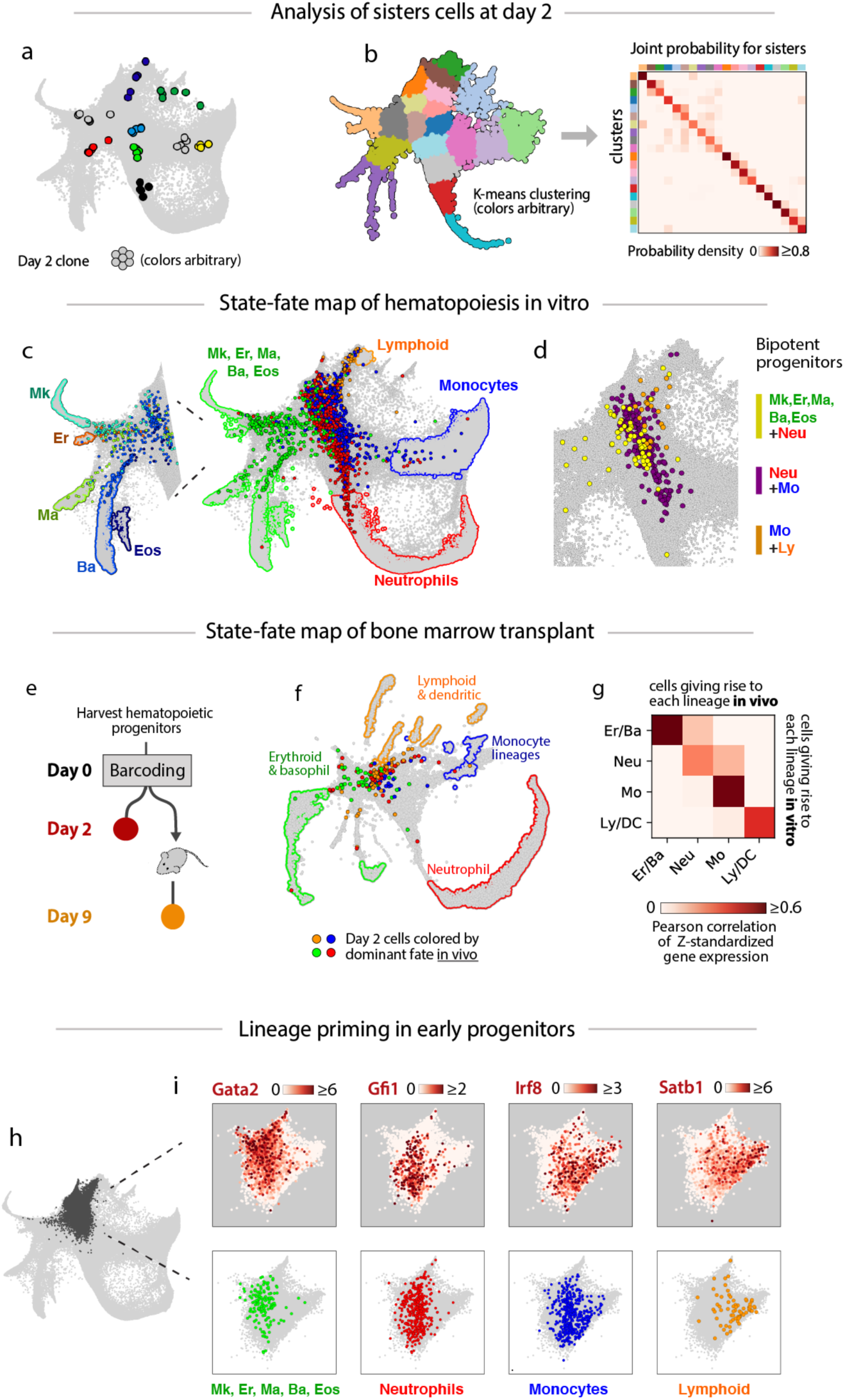
Clonal dynamics of hematopoietic progenitor culture in vitro. (a) Examples of day 2 clones; each color denotes one clone. (b) Systematic demonstration of similarity of cells within two-day clones by clustering all cells at day 2 (left) and then calculating the probability of sister cells occupying a given pair of clusters (right). (c) Linking cell states of clones across time. Day 2 cells (colored dots) are colored by the dominant fate of their mature sisters observed at a later time. Colored outlines circumscribe regions of the SPRING plot that correspond to each respective fate of day 2 cells. (d) Location of bi-potent progenitors. Colored dots are day 2 cells that have exactly two fates among their sisters at later time points. Each color represents a different combination of two lineages. (e) Experimental setup for in vivo clonal tracking experiment. (f) Clonal lineage outcomes of cells one-week post-transplantation. (g) Gene expression similarity of lineage progenitors in vitro and in vivo. Color indicates the correlation of average normalized gene expression between day 2 cells that gave rise to each lineage in vitro and in vivo respectively. (h-i) Analysis of lineage priming in early progenitors. (h) Early progenitor cells (dark gray) identified by high expression of Cd34. (i) Expression patterns of transcriptional regulators (top) juxtaposed with the distribution of progenitors for each lineage (bottom).

We visualized the cell transcriptomes using a force-directed layout of the data (SPRING plot^19^), which revealed a continuous state map spanning from multipotency to nine mature cell types that appeared in culture (Fig. 1e): erythrocytes (Er), megakaryocytes (Mk), basophils (Ba), mast cells (Ma), eosinophils (Eos), neutrophils (Neu), monocytes (Mo), dendritic cells (plasmocytoid pDC; migratory migDC) and lymphoid precursors (Ly). The cell types appearing in culture faithfully reflected the same states observed after engraftment (Fig. 1f; Supp Fig 3a), albeit with differences in their abundance (Supp Fig 3b), and a notable depletion of lymphoid cell types in vitro, such as committed B, T and NK progenitors. The clones exhibited a range of behaviors in vitro, including uni-lineage differentiation, multi-lineage branching, and stationary self-renewal of early progenitors (Fig 1g). Therefore, with this experiment we simultaneously captured both a single cell state map, and its underlying clonal relationships.

**Figure 3:**
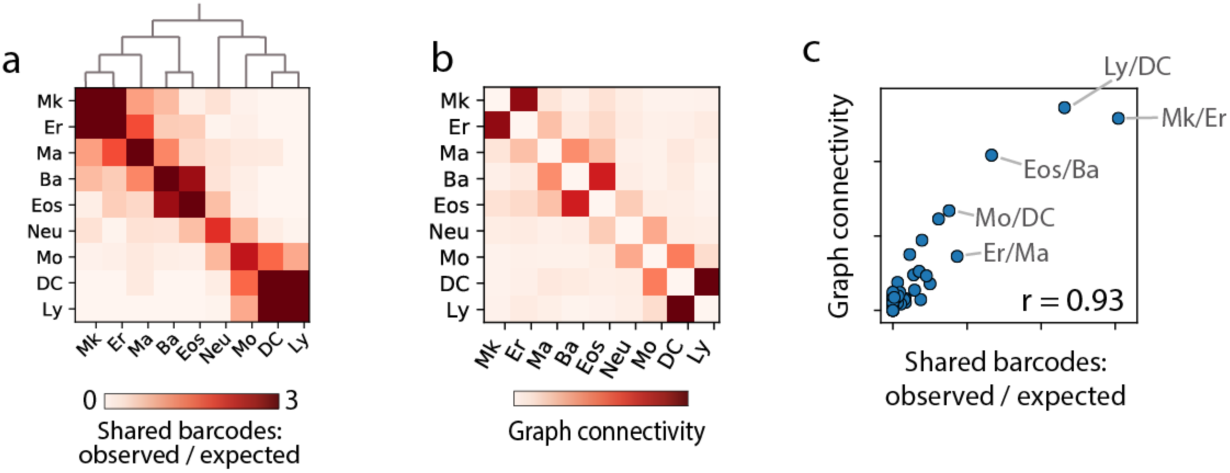
Agreement of state and lineage differentiation hierarchies. (a) Heatmap of clonal-overlap between pairs of lineages. Color indicates the observed number of shared barcodes between a paired of lineages, normalized by expectation if clonal membership is shuffled. The dendrogram (top) shows the result of iteratively joining the most coupled lineages. (b) Heatmap of state proximity for pairs of lineages. Color represents connectivity in the SPRING graph calculated by graph diffusion of cell type labels. (c) State proximity across all pairs of lineages (x-axis) correlates with their clonal relatedness (y-axis).

### Clonal dynamics identify early transcriptional fate boundaries

With LARRY it is possible to link the undifferentiated state of cells to their future fate, since sampling the same clone at an early and a late time point approximates the change that a single cell would undergo over time. The fidelity of this approximation depends critically on the similarity of sister cells at the early time point. We therefore sampled pairs of sister cells that were both profiled on day 2 and confirmed that they had similar gene expression using a range of metrics: visually, the sister cells co-localized in the SPRING plot (Fig 2a; Supp Fig 4a); they correlated well in gene expression of robustly detected genes (Supp Fig 4b; median R^2 of Z-scored, imputed expression = 0.81, compared to 0.12 for random cell pairs); and after clustering single cell data we found that sister cells overwhelmingly appeared in the same cluster across a range of clustering methods and granularities (Fig 2b; Supp Fig 4c). The sister cells were not identical, however, as they showed greater differences in gene expression than can be attributed to sampling alone (Supp Fig 4d). We tested and ruled out that sister cell similarity arises from technical co-encapsulation artifacts during scSeq (Supp Fig 4e). These tests justified examining the change in cell state within each clone across time.

**Figure 4:**
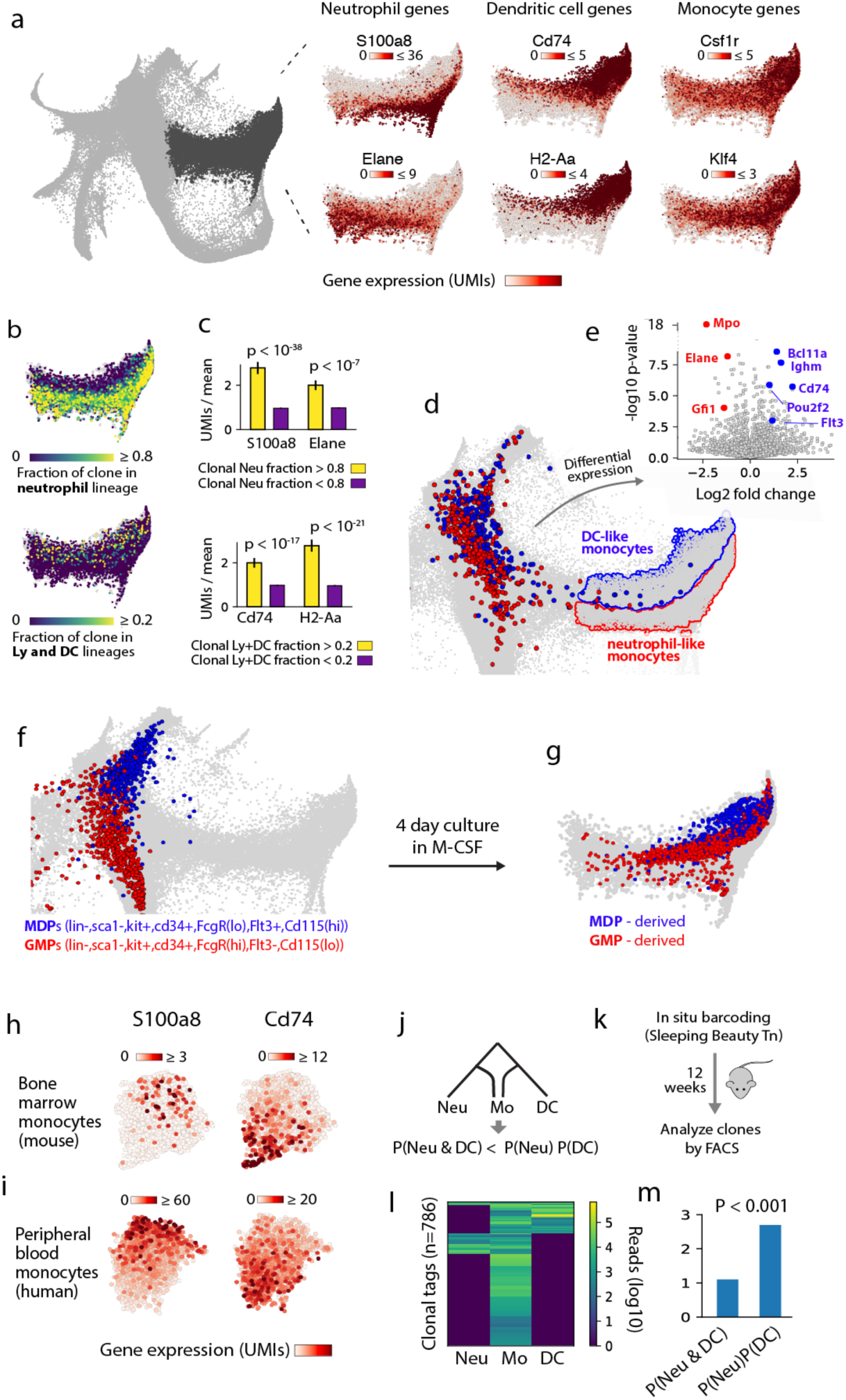
Multiple paths of monocyte differentiation. (a) Gene expression heterogeneity among differentiating monocytes, defining an axis between expression of neutrophil markers (*S100a8*, *Elane*) and DC markers (*Cd74*, *H2-Aa*), with all cells expressing monocyte markers (*Csf1r*, *Klf4*). (b-c) Clonal couplings of monocyte subsets. (b) Monocytes colored by their proportion of neutrophil sisters (top) or DC sisters (bottom). (c) Quantifying the link between gene expression and lineage history seen in panels(a,b) by differential expression of neutrophil markers among monocytes with or without significant clonal coupling to neutrophils (top), and the same for DC markers (bottom). (d) SPRING plot showing distinct transcriptional states of day 2 progenitors whose sisters differentiate into neutrophil-like (red) or DC-like (blue) monocytes, indicating two separate pathways of monocyte differentiation. (e) Volcano plot identifying differentially-expressed genes between the early (day 2) progenitors of DC-like and neutrophil-like monocytes. (f) The location of freshly sorted GMPs and MDPs projected onto the SPRING plot. (g) The projected positions of MDP-derived and GMP-derived cells after 4 days of culture in M-CSF. (h-i) A DC-to-neutrophil axis of gene expression persists in mature monocytes, as seen by SPRING plots of scSeq data from monocytes in mouse bone marrow (top) and human blood (bottom). (j-m) Clonal analysis of monocyte differentiation in unperturbed hematopoiesis. (j) Under a model of two different monocyte differentiation pathways, Neu-DC-Mo clones should be depleted relative to the null expectation. (k) Experimental schematic for barcoding mouse bone marrow in situ with clonal cell type composition assayed after a 12-week chase. (l) Heatmap showing the number of cells in each lineage (columns) detected for each lone (rows). (m) Clonal analysis shows a depletion of Mo-Neu-DC clones, consistent with two-pathway model.

We visualized the fate map of early (day 2) cells by labeling each cell according to the majority fate among its sisters on days 4 and 6 (Fig. 2c). This map revealed that already at day 2, cells separated into well-delineated domains of dominant fate potential. There were only limited areas of overlap between the progenitor domains for different lineages, apparent in the plot as salt- and-pepper mixtures of differently marked cells. Upon examination, we found that these regions represented uncommitted states, containing cells that individually gave rise to two or more clonal fates at later time points (Fig 2d).

Similar fate boundaries were also detected for cells transplanted *in vivo*, though blurred, possibly due to lower statistical power (Fig 2e,f analyzing n=104 clones that appeared both before and one week after transplantation). We noted one major difference between the in vivo and in vitro experiments in the identity of neutrophil progenitors, which was restricted to the most immature cells in the in vivo experiment, likely due to the fast rate of neutrophil differentiation and the longer chase period in vivo (7 days as opposed to 4 days in vitro). With the exception of neutrophils, however, the progenitors for each lineage mapped to their corresponding in vitro transcriptional states (Fig 2g).

We next asked how fate potential relates to underlying patterns of gene expression. Returning to the in vitro data and focusing on the earliest progenitors (Fig 2h; region identified by high expression of Cd34), we observed that the regions of bias toward each lineage overlapped closely with the expression of known hematopoietic transcription factors (Fig 2i) (e.g. *Gata2*^*^20^*^, *Gfi1*^*^21^*^, *Irf8*^*^22^*^ *and Satb1*^*^23^*^). Overall, these observations support the view functional lineage priming varies across a continuous hematopoietic progenitor landscape and covaries with the heterogenous expression of key lineage transcription factors.

### Agreement of state and lineage differentiation hierarchies

Access to state and lineage information additionally allows us to reflect on a common assumption in scSeq studies of differentiation: that lineages with transcriptionally similar differentiation pathways are clonally related. This intuition underlies all methods for hierarchy reconstruction from single cell data^3-5,7^. However, similar end states could also arise from non-overlapping clones^24^ and distant end states could share lineage through asymmetric division. In hematopoiesis, the granulocyte lineages (Ba, Eos, Ma and Neu) have similar functional properties but may have distinct ontogenies, with a recent proposal that basophils and eosinophils share a progenitor with erythrocytes and megakaryocytes, rather than neutrophils as previously thought^25,26^. To address how clones disperse on the cell state landscape, we quantified the ‘lineage coupling’ of each pair of differentiated states from their number of shared clones (Fig. 3a), as well as the similarity of cell states (Fig. 3b) using the in vitro data (see metrics in figure captions). We found that measures of state distance and clonal coupling were closely correlated across a broad range of parameter choices (r=0.93, *p <* 10^-35^; Fig 3c, Supp Fig 5a), and for different distance metrics (Supp Fig. 5b-c). The concordance of state distance to clonal coupling can also be appreciated by the agreement between the unsupervised layout of cell states (Fig. 1e) and the dendrogram of cell states based on clonal distance (Fig. 3a). Notably, both state and fate analysis confirmed the existence of a common Ba-Eos-Ma-Er-Mk progenitor in vitro. We conclude that, at least in vitro, HPC clones disperse in a hierarchical manner over the cell state continuum, in agreement with classical notions of hematopoiesis^8^.

**Figure 5:**
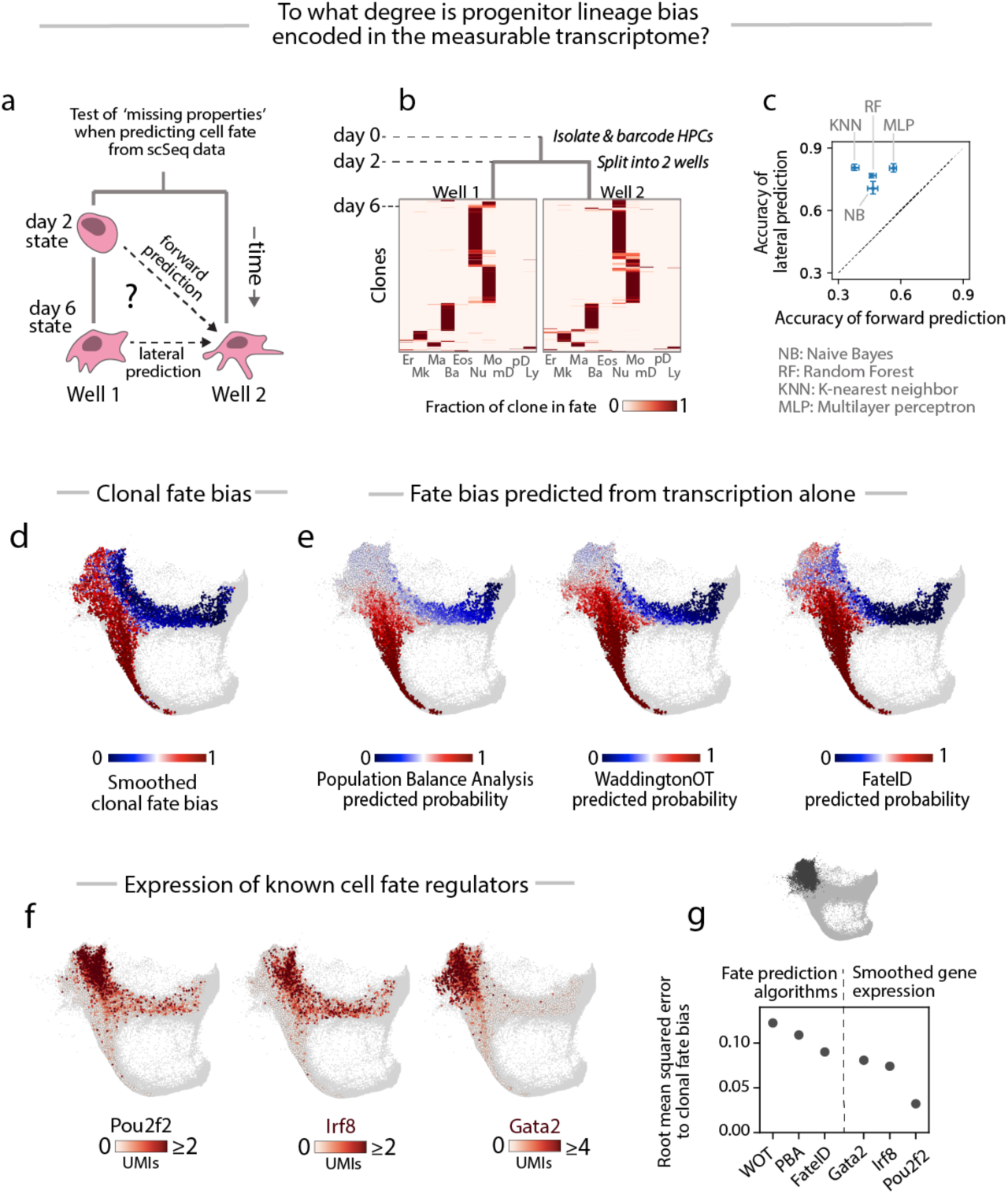
A benchmark for fate prediction in hematopoiesis. (a) Split-well experimental setup for testing whether cells have heritable properties that influence fate choice but are not detectable by scSeq. Hidden heritable properties would manifest if cell fate outcomes are better predicted by day 6 sister cell fate (lateral prediction) than by day 2 state (forward prediction). (b) Clonal fate distributions for clones that were split into different wells on day 2 and then profiled on day 6. Each row across both heatmaps is a clone; color indicates the proportion of the clone in each lineage in the respective wells. (c) Comparison of the accuracy of forward prediction (x-axis) and lateral prediction (y-axis). Data points indicate results of different machine learning methods [naïve Bayes (NB), k-nearest neighbor (KNN), random forest (RF), multilayer perceptron (MLP)], with error-bars representing 25^th^ and 75^th^ percentiles of prediction accuracy across 100 partitions of the data into training and testing sets. (d) SPRING plot showing the neutrophil/monocyte differentiation trajectory (excised from Fig. 1e), with day 2 cells colored by the smoothened ratio of neutrophil vs. monocyte fate of their sisters on days 4-6. (e) Comparison of algorithms that predict fate probability from transcriptional information alone. Color represents predicted probability of neutrophil versus monocyte differentiation. (f) Expression of cell fate regulators, showing differential expression among the earliest progenitors of neutrophil and monocyte fate, before lineage bias is recognized by the algorithms shown in (e). (g) Standard error of the algorithmically predicted fate probability compared to the observed values (left), versus smoothened gene expression (right), among early progenitor cells (dark gray region, top).

### Monocytes differentiate via multiple trajectories

In the data, monocytes appeared to form a spectrum from neutrophil-like to DC-like, expressing alternatively neutrophil elastase (Elane) and other neutrophil markers, or MHC class II components (Cd74 and H2-Aa) (Fig 4a; Supp Fig 6a). No similar overlap occurs with other lineages (Supp Fig 6b). We investigated whether this phenotypic spectrum might result from distinct differentiation trajectories of monocyte progenitors, as recently proposed^27^, which would manifest in different clonal fate biases.

**Figure 6:**
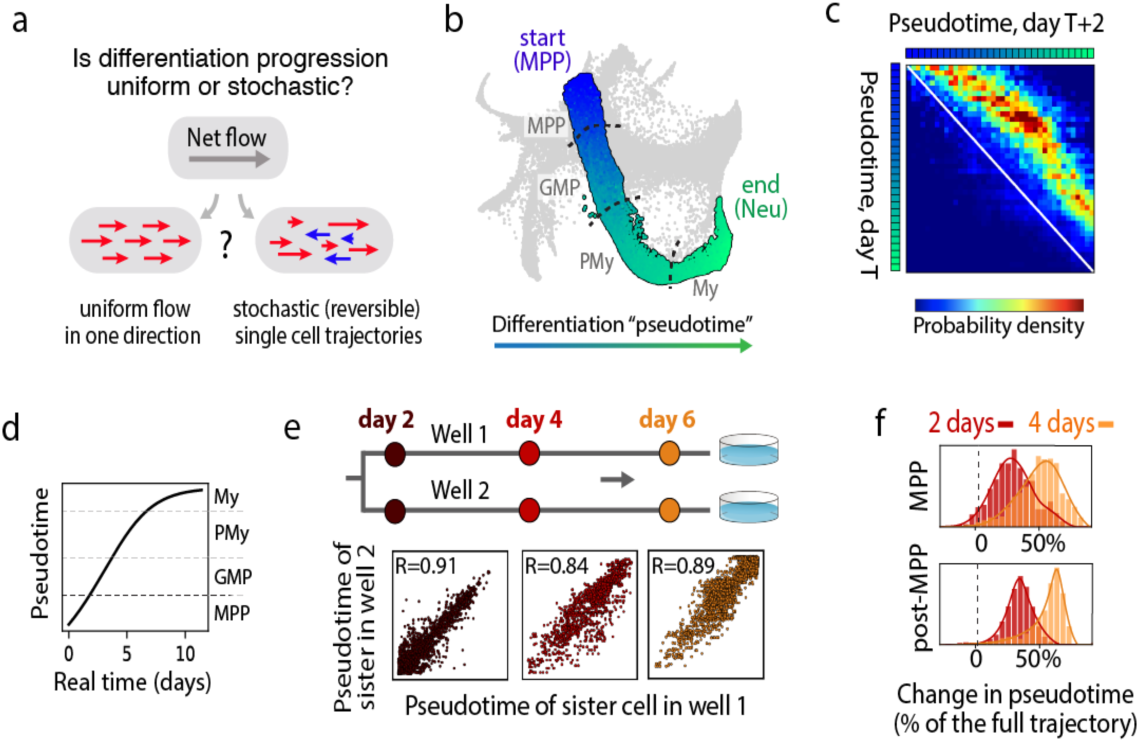
Whole-transcriptome real-time dynamics of differentiation. (a) Different alternatives for the single-cell kinetics of differentiation progression. (b) Neutrophil differentiation trajectory, colored by “pseudotime”, a measure of progression along the trajectory. Dotted lines represent the approximate boundaries in expression of marker genes associated with stages of neutrophil differentiation (PMy=promyelocyte; My=myelocyte). (c) Joint distribution of pseudotime in pairs of sister cells separated in time by 2 days reveals a consistent forward shift across the trajectory. (d) Integration of pseudotime velocity from panel(c) shows pseudotime progress as a function of real time for a typical cell. (e) Comparison of pseudotime for sister pairs that were separated on day 2 and cultured in separate wells. (f) Distribution of change in pseudotime from day 2 to day 4 (red) or day 6 (orange). Clones are stratified by their stage on day 2, with MPPs (top) showing a greater variability of pseudotime change than post-MPP stages (bottom).

To determine whether monocytes might have multiple ontogenies, we scored their clonal relatedness with mature neutrophils and DCs (Fig 4b). We found that monocytes were not uniformly coupled to either lineage: those with increased expression of neutrophilic markers shared lineage with neutrophils (Fig 4c; *p <* 10^*-*7^, Mann-Whitney U test), whereas those with DC-like gene expression shared lineage with DCs and lymphoid cells (Fig 4b,c; *p <* 10^*-*17^). We did not observe a comparable phenomenon for any other lineage in our data. Thus, monocytes appear unique in showing a phenotypic spectrum that correlated with distinct clonal histories.

The distinct clonal origins of monocytes suggested that they should arise from progenitors with different fate potentials, and possibly different gene expression profiles. To test these predictions, we classified the monocytes appearing after 4-6 days as either DC-like or neutrophil-like, and then examined their clones at the earliest time point (2 days). Indeed, the predecessors of DC-like and neutrophil-like monocytes segregated clearly by gene expression (Fig 4d,e), with respective expression of early DC markers (Flt3, Bcl11a and Cd74) or early neutrophil markers (Elane, Mpo and Gfi1). These results indicate that in our culture system, there are two different pathways of monocyte differentiation that have distinct lineage relationships and gene expression.

These results are consistent with a recent finding that immunophenotypically-defined monocyte-dendritic progenitors (MDPs) and granulocyte-monocyte progenitors (GMPs) give rise to monocytes with DC-like and neutrophil-like characteristics, respectively^27^. To test whether our observations represent the differential outputs of such populations, we performed scSeq on fresh MDPs and GMPs sorted from adult mouse bone marrow and found that they co-localized with the day 2 progenitors of DC-like and neutrophil-like monocytes (Fig 4f). Similarly, scSeq analysis of MDPs and GMPs cultured for 4 days in vitro co-localized with mature DC-like and neutrophil-like monocytes (Fig 4g). Thus, the DC-like and neutrophil-like trajectories observed here likely represent MDP and GMP pathways of monocyte differentiation, and they clarify the location of these fate-biased states in a gene expression continuum.

These results raise the question of whether the DC-like and neutrophil-like monocytes emerge through distinct clonal dynamics in unperturbed hematopoiesis. Mature bone marrow monocytes have been shown to exhibit a DC-to-neutrophil spectrum of gene expression^27^, which we also observed in classical monocytes (Fig 4h), and human peripheral blood monocytes (Fig 4i). If monocytes arise via distinct paths in steady-state conditions, we would expect an anti-correlation between neutrophil and DC relatedness among monocytes (Fig 4j). We examined the clonal co-occurrence of monocytes with DCs and neutrophils after genetically barcoding HPC clones in a non-perturbative manner using a transposase-based strategy^28^. After a 12 week chase (Fig 4k), we indeed found significantly fewer neutrophil-monocyte-DC tags than would be expected if clonal co-occurrence were independent (2.5-fold reduction; p < 0.001 by binomial test of proportion; Fig 4l,m). Overall, our results support the existence of multiple monocyte ontogenies in native hematopoiesis as well as in culture.

### A benchmark for fate prediction in hematopoiesis

In attempting to understand the physiology of hematopoietic progenitors, we and others have been interested in developing data-driven models of cell fate choice^3-5,7,26^, and have been strongly motivated by the rich data of scSeq. Such models can be used, for example, to predict cellular components driving fate choice. But the success of any predictive model – no matter how complex – requires critically that cell measurements are informative of behavior. For scSeq this basic requirement has not been testable until now. Here, with access to clonal dynamics we have a benchmark against which fate predictions from scSeq data can be tested. We set out to address three questions: (1) in principle, how well could the measured transcriptional state of a cell predict its fate? (2) how well do existing computational methods perform in making fate predictions? And (3), how accurately does scSeq reconstruct temporal dynamics? These questions are focused here on hematopoiesis but are broadly relevant to researchers applying scSeq to study dynamic processes in any system.

### The missing heritability of transcriptional states

Cell fate depends on a complex interaction between transcription, epigenetic state, cell signaling and other factors. Therefore, it is possible for two cells to appear the same from the perspective of scSeq, yet already be committed to different fates as a result of stable non-transcriptional properties. These properties, which can be thought of as ‘hidden variables’^4^, would fundamentally limit the accuracy of fate prediction from scSeq. To test whether hidden variables influence cell state, we compared the accuracy of predicting a cell’s fate from the state of its sister cell, either before the sister has differentiated (‘forward prediction’), or after the sister has differentiated (‘lateral prediction’; Fig 5a).

If there were no hidden variables, then the mutual information between sister cells could only decrease as time passes since they divided, an intuition formalized in information theory as the *Data Processing Inequality*. Thus, when predicting the fate of a cell from the state of its sister, nothing would be gained by waiting for the sister to differentiate. On the other hand, if there are stable properties that influence cell fate but are hidden from the transcriptome, then the apparent mutual information between sisters could increase over time as these properties manifest in their choice of cell fate. These two scenarios can be distinguished by comparing the accuracy of forward prediction to that of lateral prediction (Fig 5a). If lateral prediction is more accurate, then there must be stable properties influencing cell fate that are hidden from the transcriptome (see Methods for a mathematical argument).

We have so far obtained data for forward prediction (Fig 2c), but not from a late state. To assess the accuracy of lateral prediction, we barcoded Lin-Kit+ Sca1+ HPCs (LSK cells), split them into two wells after 2 days, and then differentiated them in the separate wells for 4 further days. In total, we identified n=502 barcodes appearing in both wells, consisting of 6351 cells that evenly covered the cell state map (Supp Fig 7a). These barcodes seen in both wells correspond to the progeny of sister cells, allowing the fate of sisters to be compared.

Remarkably, we observed a high rate of concordance in the fates of clonally related cells in different wells (Fig 5b): clones in separate wells produced identical combinations of lineages 70% of the time, compared to 26% by chance. This concordance held for clones of all potencies, and is consistent with recent observations of clonal fate restriction among HPCs^10^. To compare the accuracy of forward prediction versus lateral prediction, we applied a panel of machine learning algorithms to guess the dominant fate of a clone using either the transcriptomes of its day 2 sisters, or the transcriptomes of its day 6 sisters, using the fates and transcriptomes of other cells as a training set. We found that lateral prediction was more informative for all algorithms tested (Fig 5c), with the most accurate algorithm – a neural network – showing 26% less accuracy for forward prediction compared to lateral prediction.

These results imply that some heritable properties of cell physiology influence cell fate but are not evident in our scSeq measurements. Missing information of this type is akin to the notion of “missing heritability” in population genetics^29^. We cannot tell whether information on cell fate is restricted simply because scSeq data is noisy, or because cell fate depends on cellular properties that are not reflected in the transcriptome, such as chromatin state, protein abundances, cell organization, or the microenvironment. Yet we may conclude that, at least for in vitro hematopoiesis, transcriptional models could in principle be predictive of cell fate, but not entirely so.

### scSeq-based models do not fully predict fate choice

We next asked how well existing computational models, using only scSeq data, predict cell fate probabilities. We tested three recent approaches, Population Balance Analysis (PBA)^4^, WaddingtonOT (WOT)^5^, and FateID^7^ for their ability to predict the fate of a cell choosing between neutrophil and monocyte fates. These cells occupy a landscape that is densely sampled by cells and clones (96,373 cells, 4279 clones) and so provides a good test case. We calculated for each cell at day 2 *in vitro* the fraction of its clonal relatives that became a neutrophil or a monocyte (Fig 5d), and then attempted to predict this fraction from transcriptome snapshots alone (Fig 5e). We found that all three methods were similarly accurate with a clear advantage to FateID (Pearson correlation to empirical fate bias of r=0.68, 0.71, 0.78 for WOT, PBA and FateID respectively at the optimal parameter setting for each method; Supp Fig 7b). However, neutrophil and monocyte lineage bias occurs earlier than predicted by any of the algorithms, in cells still expressing markers of MPPs that do not show a discrete bifurcation in gene expression state.

Overall, these results show that in the absence of lineage information, computational methods for scSeq analysis may mis-identify fate decision boundaries. It is therefore significant that early lineage bias can be distinguished based on transcriptional information including the expression of known fate regulators such as Gata2 and Irf8 (Fig 5f). Indeed, when restricting to early progenitors (Cd34-high cells) the imputed expression of just a single gene, *Pou2f2*, provided a superior prediction of cell fate than algorithmic predictions (Fig 5g). The selection of the correct genes to use for prediction, however, required clonal information. We obtained similar results when we repeated the analysis on the full dataset, including all lineages (Supp Fig 7c-e). These results provide a framework for comparing computational models of differentiation, and may serve as a useful benchmark for improving them.

### Benchmarking the noise in differentiation dynamics

As cells differentiate, they progress through a sequence of gene expression intermediates that chart a trajectory in high dimensional gene expression space. A common goal of scSeq is to determine a typical gene expression trajectory by defining a ‘pseudotime’ coordinate that orders the transcriptomes along the trajectory^30^. At present, it is unknown how single cells traverse these trajectories, including whether they progress at different rates or even reverse their dynamics^4^ (Fig. 6a). Focusing on neutrophil differentiation as a test case, we asked how well ‘pseudotime’ describes the kinetics of differentiation as revealed by clonal tracking.

By ordering of cells from MPPs to GMPs, to promyelocytes (PMy), to myelocytes (My) (n=63,149 cells Fig. 6b; Supp Fig 8a,b), we could compare the pseudotemporal progression of clones sampled at two consecutive days (Fig. 6c). This analysis showed a strikingly consistent forward velocity along differentiation pseudotime. By integrating the velocity across the trajectory, we were able to calculate pseudotime progression as a function of real time for a typical cell (Fig. 6d). The time for an MPP to differentiate into a myelocyte was 10 days, consistent with prior literature^31^. Pseudotime analysis of sister cells differentiated in separate wells also showed a remarkable consistency in the pace of differentiation. On day 2, shortly after cell division, the pseudotime of sister cells had a correlation of R=0.91 (Fig 6e). Four days later, the pseudotime correlation for sister cells in separate wells remained almost as high (R=0.89; Fig 6e). However, pseudotime velocity was not uniform: maximal variability occurred among MPPs and decreased for subsequent stages (Figs. 6f). This could be explained by cells remaining in the MPP state for a variable duration before initiating neutrophil differentiation. These results support the use of pseudotime methods for mapping differentiation progression.

## DISCUSSION

We describe here a new scSeq-compatible lentiviral lineage tracing tool that, combined with a clone-splitting experimental design, allows tracking cells simultaneously from multiple initial conditions, without needing to target each specific progenitor state. In doing so, it delineates fate boundaries in an unbiased manner, avoiding boundaries imposed by a particular choice of reporter gene or by cell sorting criteria. The method additionally offers an advantage over scSeq analysis alone by providing definitive temporal links between different transcriptional states. These advantages allowed us explore questions that could not be addressed with conventional lineage tracing or scSeq.

In hematopoiesis, clonal tracing studies have shown evidence of functional lineage priming of multipotent progenitor cells (MPPs)^32^. Separately, scSeq studies have shown that MPPs are transcriptionally heterogeneous, existing along a continuum with transcriptional priming toward different lineages^26^. Here we unify these views by locating functionally primed cells on a gene expression continuum, showing that regions of functional lineage priming in MPPs associate with low-level expression of lineage-affiliated transcription factors. From these biased states, cells differentiate via a structured fate hierarchy that differs from classical tree-like depictions of hematopoiesis in its clonal structure and confirms recently-established ontogenies of granulocytes^25,33^ and monocytes^27^.

Our study also allowed us to critically evaluate approaches for inferring cell dynamics from static scSeq data. We showed that fate biases appear much earlier than predicted by several methods tested and, significantly, that fate prediction from scSeq data is limited by missing information on the state of the cell. Such missing information, which represents heritable properties of cell physiology that eventually manifest in the fate of sister cells, cannot be deciphered from scSeq data alone, at least given the sampling constraints of current scSeq methods. Despite these limits, we showed that the overall structure of a differentiation hierarchy, and the ordering of events along one branch, can be predicted well by scSeq alone. Our results therefore offer much-needed benchmarks for testing scSeq analytical algorithms.

It is important to emphasize several limitations of the method. First, it is currently restricted to culture or transplantation assays, and it requires sister cells to be separated. This limitation disrupts spatial organization and so restricts potential applications. Additionally, because cells must divide at least once, the method cannot study processes faster than one cell cycle. Yet within these constraints, the method opens up the possibility of perturbation approaches that directly compare the fates of sister cells treated differently yet closely matched in their initial state, and offers benchmarks for *in vivo* experimental and computational scSeq and lineage tracing. Coupling cell state and fate readouts in different tissues will deepen our understanding of stem cell behaviors in tissue development and homeostatic physiology.

## FUNDING

A.M.K. and C.W. were supported by NIH grants R33CA212697-01 and 1R01HL14102-01, the Harvard Stem Cell Institute Blood Program Pilot Grant DP-0174-18-00, and the Chan-Zuckerberg Initiative grant 2018-182714. A.R-F was supported by the Merck Fellowship of the Life Sciences Research Foundation, a EMBO Long-Term Fellowship (ALTF 675-2015), an American Society of Hematology Scholar Award and a Leukemia & Lymphoma Society Career Development Program Award (3391-19). F.D.C was supported by NIH grants HL128850-01A1 and P01HL13147. F.D.C. is a scholar of the Howard Hughes Medical Institute and the Leukemia and Lymphoma Society.

## Competing interests statement

AMK is a founder of 1Cell-Bio, Ltd.

## Acknowledgements

We thank the Single Cell Core Facility at Harvard Medical School for inDrop reagents; the Bauer Core Facility for sequencing; Bertie Gottgens for discussions and comments on the manuscript; and Kyogo Kawaguchi for mentorship in experiments and analysis.

## Author contributions

All authors designed the experiments. C.W and A.R-F performed the experiments. C.W. carried out the computational analysis. C.W. and A.M.K. wrote the paper. A.M.K. and F.D.C jointly supervised the work.

## Data availability

Data and jupyter notebooks for select analyses are available at https://bit.ly/2z2F8jX

**Table 1:**
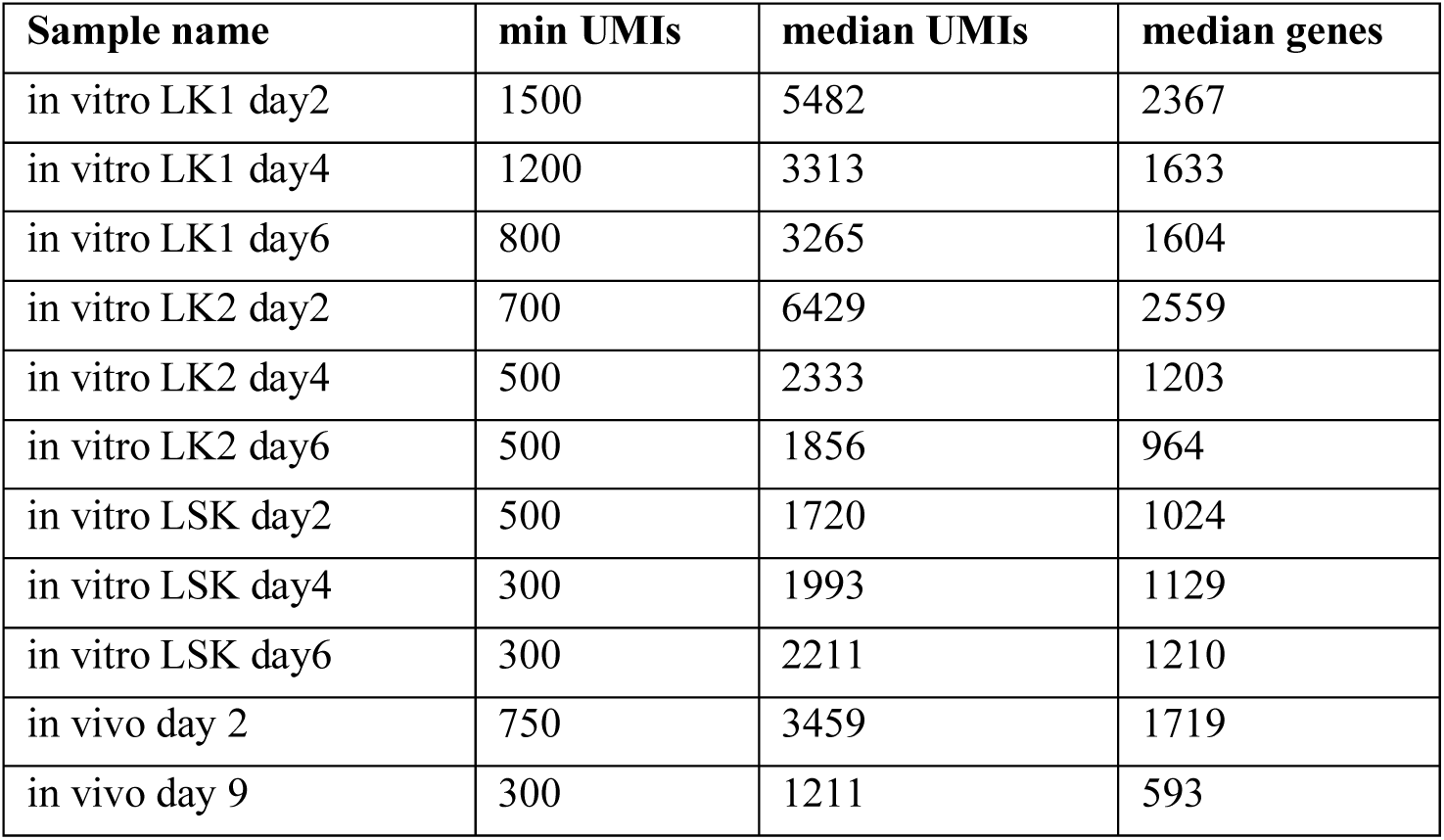
UMI thresholds for cell filtering

| Sample name | min UMIs | median UMIs | median genes |
| --- | --- | --- | --- |
| in vitro LK1 day2 | 1500 | 5482 | 2367 |
| in vitro LK1 day4 | 1200 | 3313 | 1633 |
| in vitro LK1 day6 | 800 | 3265 | 1604 |
| in vitro LK2 day2 | 700 | 6429 | 2559 |
| in vitro LK2 day4 | 500 | 2333 | 1203 |
| in vitro LK2 day6 | 500 | 1856 | 964 |
| in vitro LSK day2 | 500 | 1720 | 1024 |
| in vitro LSK day4 | 300 | 1993 | 1129 |
| in vitro LSK day6 | 300 | 2211 | 1210 |
| in vivo day 2 | 750 | 3459 | 1719 |
| in vivo day 9 | 300 | 1211 | 593 |

**Table 2:**
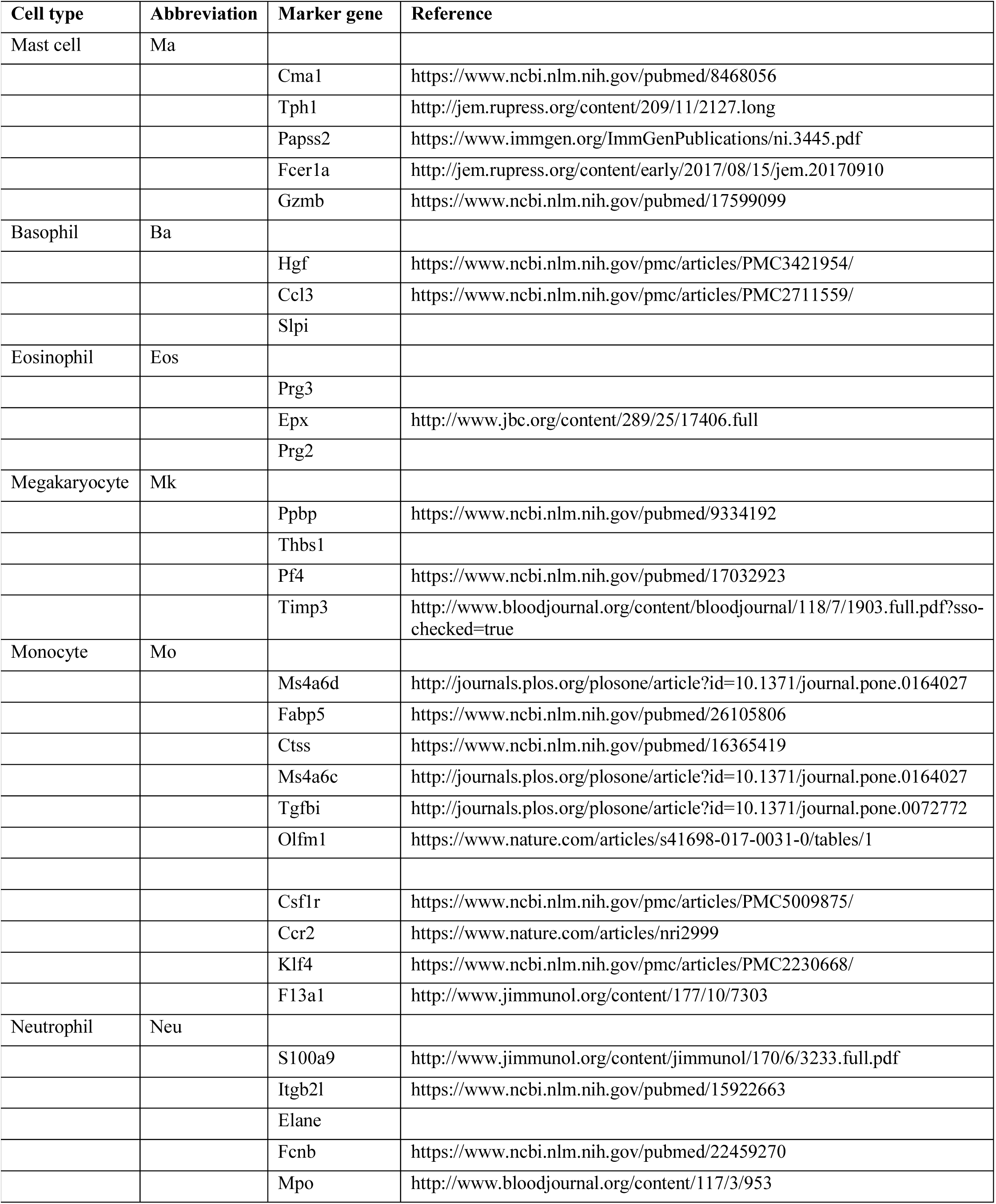
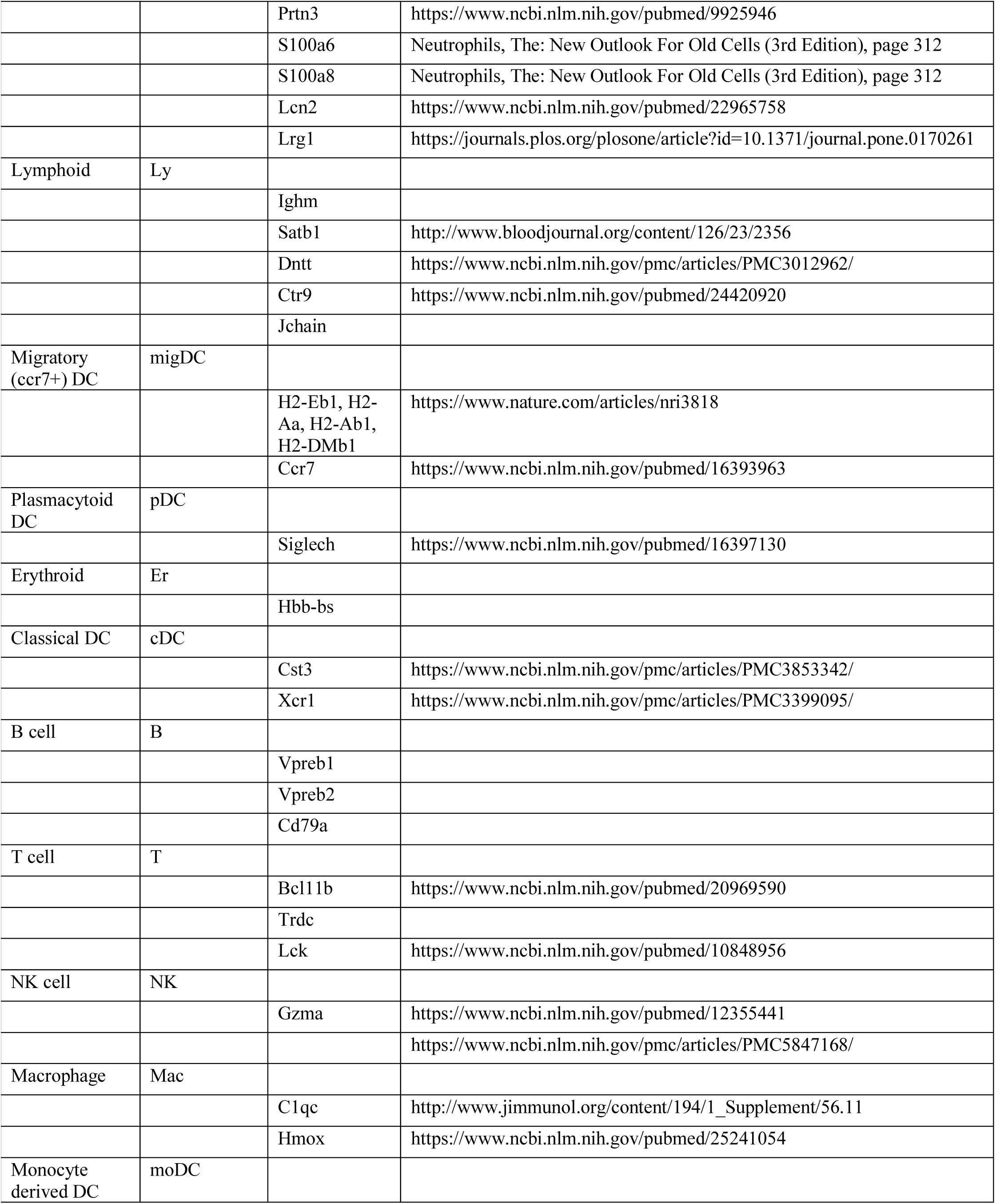
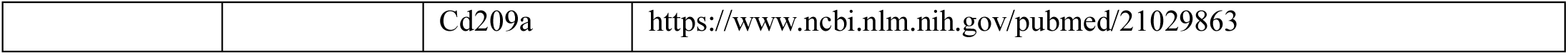
Marker genes used for cell type annotation

## Cell culture and lentiviral barcoding

### Barcoded lentiviral library synthesis and amplification

The pLARRY vector was constructed by DNA synthesis and Gateway cloning (Vectorbuilder) using a protocol loosely adapted from (Naik, Schumacher et al. 2014) and (Gerrits, Dykstra et al. 2010). The barcoded linker was created by annealing two DNA primers (forward, 5′-CCC CGGATCCAGACATNNNNCTNNNNACNNNNTCNNNNGTNNNNTGNNNNCANNNNCATATGAGCAATCCCCACCCTCCCACCTAC-3′; reverse, 5′-GTAGGTGGGAGGGTGGGGATTGCT-3′; IDT DNA). N was a hand mix of 25% A, 25% C, 25% T and 25% G. Primers (10 pmoles of each) were mixed in 50 µl 1× NEB buffer 4 (New England Biolabs). After heating the mixture for 5 minutes at 95°C, the primers were allowed to anneal down to 37°C gradually decreasing temperature (0.5°C/minute). Then, 1U of Klenow DNA polymerase (3’-5’ Exonuclease mutant) and 50 nmoles of dNTPs was added to the mixture and incubated for 2 hours at 37°C. After Klenow inactivation for 20 minutes, the barcoded linker was then digested with a mixture of NdeI and BamHI (New England Biolabs) and ligated into the *NdeI*-*Bam*HI site of the pLARRY vector at 3:1 ratio. The resulting ligation mix was purified and transformed into 10-beta electroporation ultracompetent *Escherichia coli* cells (New England Biolabs) and grown overnight on LB plates supplemented with 50 µg/mL ampicillin (Sigma-Aldrich). From 8 plates, ∼0.5-1×10^6^ colonies were pooled by flushing plates with LB supplemented with 50 µg/mL ampicillin. After 6h of culture, plasmid DNA was extracted with a Maxiprep endotoxin-free kit (Macherey-Nagel). The pLARRY vector map and plasmid, as well as a sample of the library will be made available through Addgene before publication.

### Lentivirus production and barcode labeling

Barcoded GFP plasmid and third generation lentivirus components were transfected into HEK293T cells using the Lipofectamine 3000 kit (Thermofisher). Lentivirus was harvested every 12 hours for 72 hours and concentrated using ultra-centrifugation at 80,000g. HEK cells were grown in DMEM with 10% fetal bovine serum (FBS) and 1% PenStrep but switched to OptiMEM (Thermofisher) with 10% FBS for transfection and lentivirus harvest. Hematopoietic progenitor cells (HPCs) were transduced using spin infection (800g for 90 minutes) in virus concentrate with DAEA-dextran hydrochloride (Sigma).

### Cell isolation for state-fate experiments

After euthanasia, bone marrow from femur, tibia, pelvis and sternum was isolated by crushing with pestle and mortar to obtain all cells. Collected bone marrow cells were filtered through a 40-µm strainer and washed in cold ‘Easy Sep’ buffer (PBS; 2% fetal bovine serum (FBS); 1?mM EDTA; Pen/Strep). Red blood cells were then lysed using ammonium chloride (STEMCELL) and mature lineage cells were depleted magnetically using the EasySep Lineage Depletion Kit (STEMCELL). The resulting Lin-fraction was stained for Kit (CD117-PE, clone 2B8, Biolegend), Sca-1 (Ly6a-FITC, clone D7, Biolegend) and lineage markers (antibody mix from the same EasySep kit), and Lin-Kit+Sca-1+ (LSK) or Lin-Kit+ (LK) cells were isolated by flow activated cell sorting (FACS) on a Sony SH800 with a 130uM nozzle.

### Cell culture for state-fate experiment in vitro

In three separate in vitro experiments, sorted LK (two experiments) or LSK cells (one experiment) were barcoded as described above and then plated in round-bottom 96-well plates in media designed to support pan-myeloid differentiation, consisting of StemSpan media (STEMCELL), Pen/Strep, IL-3 (20ng/mL), FLT3-L (50ng/mL), IL-11 (50ng/mL), IL-5 (10ng/mL), EPO (3U/mL), TPO (50ng/mL) (Peprotech), and mSCF (50ng/mL) and IL-6 (10ng/mL) (R&D Systems). The number of cells plated varied from 5,000 – 10,000. This range in the number of cells was chosen in particular to maximize the yield of multi-cell clones across multiple time points. After two days in culture, cells were split evenly, with one half being sequenced immediately and the other half re-plated in two separate culture wells. After two more days in culture, cells were again split, with 30% taken for inDrops and the remainder re-plated in 6 wells. After two further days in culture, GFP+ cells were sorted using FACS and profiled using inDrops. For the latter pair of inDrops experiments, cells were dissociated from the well by 5-minute treatment with 2.5% Trypsin (Thermofisher).

### Cell culture for state-fate experiment in vivo

Cell isolation, barcoding and culturing for 2 days was carried out as in the in vitro experiment (above), except that LSK and LK cells were this time combined into one sample, with LSK cells enriched to 33% of the total (rather than their natural 10% proportion in bone marrow). After 2 days in culture, half of the cells were analyzed by inDrops scSeq, and the other half were transplanted retro-orbitally into C57BL/6J sub-lethally irradiated host mice (500 cGy) as described in (Perie, Duffy et al. 2015). After one week, the transplanted cells were harvested from total bone marrow and spleen. We then performed red blood cell lysis, stained with Ly6G-PE (clone 1A8, ThermoFisher) to detect neutrophils and sorted into GFP+ neutrophils and GFP+ non-neutrophils on the Sony SH800 sorter using a 70uM nozzle. All GFP+ non-neutrophils and 30% of GFP+ neutrophils were then profiled using inDrops.

### Techniques for preventing cell loss

At all steps during cell culture, extreme care was taken to prevent loss of cells, since the number of barcodes shared between time points is highly dependent on maximizing the yield of profiled GFP+ cells from the total that are in culture. To minimize cell loss: all spins were performed at 500g for 5 minutes in 1.5mL tubes in a swinging-bucket centrifuge; any transfer of cells out of a well involved several washes of the well with PBS; and washing steps, including those for inDrops and re-plating, were performed using PBS with 0.5% BSA. Since measuring cell density with a hemocytometer involves loss of cells, all experiments were accompanied by an identical ‘decoy’ copy of the experiment carried out in parallel that was used to assess cell density, thus leaving the ‘real’ experimental sample unperturbed by cell counting.

## Single-cell encapsulation and data pre-processing

### Single-cell encapsulation and library preparation for sequencing

For single-cell RNA sequencing (scSeq), we used inDrops (Klein, Mazutis et al. 2015) following the protocol described in (Plasschaert, Zilionis et al. 2018), with a modification to allow targeted sequencing of the LARRY barcode. In brief, single cells were encapsulated into 3-nl droplets with hydrogel beads carrying barcoding reverse transcription primers. After reverse transcription in droplets, the emulsion was broken and the bulk material was taken through: (i) second strand synthesis; (ii) linear amplification by in vitro transcription (IVT); (iii) amplified RNA fragmentation; (iv) reverse transcription; (v) PCR. To specifically amplify barcode-containing GFP transcripts, we followed a protocol similar to (Wagner, Weinreb et al. 2018). We split the amplified RNA fraction (after step (ii)) and used one half for standard library preparation and the other half for targeted lineage barcode enrichment. To target the barcode, we modified the subsequent steps of library prep by (i) skipping RNA fragmentation; (ii) priming reverse transcription using a transcript specific primer at 10mM (TGAGCAAAGACCCCAACGAG); introducing an extra PCR step using a targeted primer (8 cycles using Phusion 2X master mix; Thermofisher; primer sequence = TCG TCG GCA GCG TCA GAT GTG TAT AAG AGA CAG NNN Ntaa ccg ttg cta gga gag acc atat) and 1.2X bead purification (Agencourt AMPure XP). Targeted and non-targeted final libraries were pooled at equimolar ratios for initial sequencing. Subsequent re-sequencing was performed on non-targeted libraries alone.

### Read alignment and cell filtering

FASTQ sequence files were demultiplexed and aligned to the GRCm38 mouse reference genome using the inDrops pipeline (https://github.com/indrops/indrops), generating cell-by-gene counts tables for each experiment and condition. Cells were filtered to include only abundant barcodes on the basis of visual inspection of the histograms of total transcripts per cell (see Table 1 for UMI thresholds for each sample, as well as statistics reporting the median UMIs and median genes detected for each cell). The data were further filtered to eliminate putatively stressed or dying cells, defined by having >20% of transcripts coming from mitochondrial genes. We then applied the SCRUBLET algorithm (https://github.com/AllonKleinLab/scrublet; Wolock et al., Cell Systems 2018) to identify putative doublet cells, i.e. cell barcodes that likely represented the combination of two or more actual cells. SCRUBLET produces a ‘doublet score’ for each cell, and these scores were thresholded based on manual inspection, being set in each case to eliminate apparent doublets – defined by co-expression of mature marker genes for different lineages – while retaining the maximum number of apparent non-doublets. Doublet filtering eliminated ∼10% of cells per dataset, in accordance with an experimentally estimated doublet rate of 5-10%. Cells within each experiment were then normalized to have the same total number of transcripts for all subsequent analyses.

### Calling of lineage barcodes

To call lineage barcodes, we began with an intermediate output of the indrops pipeline: a list of reads with annotated cell barcode and unique molecular identifier (UMI). From this list, we extracted all (Cell-BC, UMI, lineage-BC) triples that were supported by at least 10 reads, collapsed all Lineage-BC’s within a hamming distance of 3 using a graph-connected-components based algorithm, and carried forward the (Cell-BC, Lineage-BC) pairs supported by at least 3 UMIs. To call clones, we then applied a set of rigorous filtering steps: (i) Cells with the exact same set of barcodes were classified as clones; (ii) Pairs of cells in separate sequencing libraries with the same Cell-BC and Lineage-BC were discarded, since statistically these could only arise from instability of the droplet emulsion; (iii) Clones that were statistically over-abundant within a single sequencing library compared to their frequency in other libraries of the same sample were discarded, since these could also only arise through emulsion instability. These steps have been implemented in a pipeline available online: https://github.com/AllonKleinLab/LARRY.

## Data visualization and cell type annotation

### Definition of terms: “PCA-batch-correction” and “graph-smoothing”

Here we define terms that will be used repeatedly in the following methods sections. PCA-batch-correction is a method for co-embedding separate scSeq datasets in a way that minimizes their global or ‘batch’ differences. One dataset is chosen as a ‘reference’ and used to define a set of principal components that define the embedding space. Both datasets are embedded in this principal component space and then carried forward for subsequent analysis.

We use “graph-smoothing” to refer to a tunable method for local averaging in a single-cell dataset, which we adapted from (Tusi, Wolock et al. 2018). The input is a graph in which cells are nodes and edges represent similarity between cells, as well as a vector of values for each cell which is to be smoothed. At each smoothing iteration, cells update their value to be (1 – *β*)*V*_*n*_ *+ βV*_*S*_ where *V*_*n*_ is the average value of their graph neighbors and *V*_*s*_ is their own value. This operation is repeated for *N* iterations. In the following Methods sections, we use *β* = 0.1 and report the value of *N* when smoothing is invoked. Mild graph-smoothing can be interpreted as a form of imputation – allowing each cell to ‘learn’ an imputed gene expression state from the cells in its immediate vicinity. More extreme graph smoothing can be used to define a global graph coordinate based on the initial variation in the value being smoothed.

### Generation of SPRING plot layouts (Fig 1c,f)

We used SPRING (Weinreb, Wolock et al. 2018) for single-cell data visualization. For all SPRING plots shown, we began with total-counts-normalized gene expression data, filtered for highly variable genes using the SPRING gene filtering function (“filter_genes” from https://github.com/AllonKleinLab/SPRING_dev/blob/master/data_prep/spring_helper.py using parameters (85,3, 3)), and further filtered to exclude cell cycle correlated genes – defined as those with correlation R>0.1 to the gene signature defined by Ube2c, Hmgb2, Hmgn2, Tuba1b, Ccnb1, Tubb5, Top2a, and Tubb4b.

For the in vitro data (Fig 1c), we embedded cells in 50-dimensional PC space, down-sampled to 40,000 cells and imported them into SPRING dynamic mode as a k-nearest-neighbor (knn) graph with (k=4). The graph was allowed to relax in SPRING. Coordinates for the remaining ∼90,000 cells were learned by allowing each cell to choose its 40 nearest neighbors and then take on the average position of the subset of neighbors that were among the original 40,000.

For the in vivo state-fate data (Fig 1f), we aimed to generate a SPRING plot that was roughly aligned with the in vitro data. To that end, we first applied PCA-batch-correction to map the day 2 portion of the in vivo dataset against the in vitro data, and then assigned each in vivo cell the average position of its 50 nearest neighbors from the in vitro data. We then applied PCA-batch-correction to map the day 2 in vivo data against the day 9 in vivo data, constructed a knn graph with k=4, and allowed the graph to relax in SPRING, with the day 2 cells fixed to the positions that they learned from the in vitro data.

### Annotation of cell types (Fig 1e,f)

Mature cell types from each dataset were identified semi-manually, by identifying cells with high expression of lineage-specific genes (Table 2) and then refining this initial selection manually in SPRING. During manual refinement, we aimed to: (i) Include cells in an annotation if they are surrounded by other annotated cells in the SPRING plot; (ii) Exclude cells from an annotation if they are isolated from the other annotated cells in the SPRING plot; (iii) Trim the margin of an annotation to exclude cells that are near branch points in the SPRING plot and may not be fully committed. We took care to ensure that the manual refinement steps did not affect the conclusions of the paper by performing them ‘blindly’ – before any subsequent analyses were done – and not changing them from that point onward. In any case, because mature hematopoietic cell states are readily distinguishable in scSeq data, changes to the annotations would likely only affect the amount of clonal data available for analysis, rather than the qualitive features of those data. We have posted annotations for all cells online (see Data Availability).

## Sister cell similarity on day 2

### Clustering analysis of sister cell similarity (Fig 2b and Supp Fig 4c)

To assess the similarity of sister cells on day 2, we clustered day 2 cells from the in vitro dataset and computed the fraction of sister cell pairs that appear in each pair of clusters. Clustering was performed using either K-means applied to the SPRING coordinate (Fig 2b; K=20 clusters) or spectral clustering applied to PCA coordinates (Supp Fig 4c; K=20 and K=40). For each of these clusterings, we computed a probability matrix *P*, defined by:

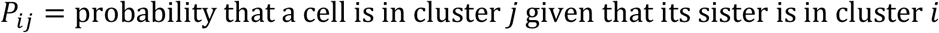

### Gene expression correlation analysis of sister cell similarity (Supp Fig 4b)

We computed the correlation of gene expression for pairs of sisters in the in vitro state-fate dataset. This analysis was restricted to highly variable genes (n=3844 genes). Gene expression values were smoothed/imputed using N=20 iterations of graph smoothing, and normalized to have the same mean and standard deviation across cells. Supp Fig 4b shows the distribution of correlation values for clonal sister pairs from day 2 versus random pairs of cells from day 2.

### Principal components analysis of sister cell similarity (Supp Fig 4d,e)

To further assess the similarity of sister cells on day 2, we computed the PCA distance between pairs of sister cells. PCA coordinates were computed from day 2 in vitro transcriptomes, total counts normalized and filtered for highly variable genes using the SPRING gene filtering function (“filter_genes” from https://github.com/AllonKleinLab/SPRING_dev/blob/master/data_prep/spring_helper.py using parameters (90,3, 3)). Supp Fig 4d shows the distribution of distance values for clonal sister pairs from day 2 versus random pairs of cells from day 2. To establish a reasonable lower bound for these distances, we also plot distances for computationally-generated statistically-identical copies of cells, as described in the next section. PCA distance was also used to compare the similarity of sisters from the same library versus distinct libraries (Supp Fig 4e), with the goal of ascertaining the contribution of gel-gel doublets to the observed sister cell similarity.

### Simulation of expected transcriptome similarity for identical cells (Supp Fig 4d)

Single-cell sequencing using inDrops only captures ∼5% of transcripts in a cell, and this down-sampling could generate the appearance of difference between cells that are biologically identical. To understand the magnitude of this sampling-induced difference, and thus establish a reasonable lower bound for sister cell similarity, we simulated pairs of identical single-cell transcriptomes as follows:

1. Pick a random cell from day 2 and identify its 20 nearest neighbors
2. Sum the raw counts for each gene across these 20 cells
3. Down-sample the counts by 20-fold. Do this twice independently to generate a pair of simulated identical cells.

## Analysis of clonal lineage couplings in vitro

### Observed/Expected clonal coupling score (Fig 3a)

We computed an observed/expected ratio of shared barcodes for each pair of lineages in the in vitro state-fate dataset. A barcode is considered shared if it appears in at least one cell from both lineages. From the observed shared barcode matrix *O*_*ij*_, we derived an expected shared barcode matrix *E*_*ij*_ under the assumption of no lineage couplings, as follows:

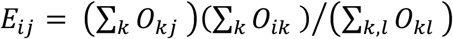

To avoid any artifacts from particularly large or atypical clones, we re-computed these matrices 500 times, each time using a random 30% sample of clones. The lineage coupling scores shown in Fig 3a represent the median *O*_*ij*_*/E*_*ij*_ from these 500 randomized trials.

## Graph connectivity score (Fig 3c)

Graph connectivity between each pair of lineages in the in vitro state-fate dataset was computed by smoothing an indicator vector centered on each lineage, as follows: For each lineage, *i*, we initialized an indicator vector *X*_*i*_ defined by:

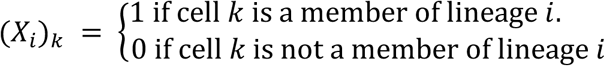

We smoothed the indicator vector using graph smoothing with N iterations, generating a smoothened vector *S*_*i*_ (N=50 iterations were used for Fig 3c and N=20 and 100 were used for Supp Fig 5a). The graph connectivity between lineages *i* and *j* was then calculated as the mean value of *S*_*i*_ among the cells of lineage *j*. Lineage membership was determined according to the annotation described above (see section “Annotation of cell types”).

## Analysis of ‘hidden properties’ that influence fate choice

We used our combined state and lineage data to ask whether there are hidden properties, such as chromatin state or protein abundance, that influence cell fate but are not evident in the measurable transcriptome. In the language of probability, these properties would cause the trajectories of cells through gene expression space to be non-Markovian (Weinreb, Wolock et al. 2018). We therefore probed our data for evidence of non-Markovian dynamics. Below, we present the theoretical justification for our approach, and then describe its practical application to data.

### Theoretical approach for analysis of hidden properties

Our approach to detect hidden properties is motivated by the data processing inequality (DPI), which formalizes the idea that in a Markov chain, information about the starting state can decrease over time but never increase. Formally, the DPI states that in a Markovian sequence of random variables *X* → *Y* → *Z*, the following inequality holds: *MI*(*X*, *Y*) *≥ MI*(*X*, *Z*), where *MI* denotes mutual information.

To apply the DPI in the context of our experiment, we consider four random variables that represent the transcriptomes of two starting sister cells (“Mother 1” and “Mother 2”) and their differentiated progeny at a later time point (“Daughter 1”, “Daughter 2”) (Supp Fig 9). If cell differentiation were Markovian with respect to scSeq measurements, then these four variables would form a Markov random field with respect to the graphical model shown in Supp Fig 9, meaning that any pair of non-adjacent nodes are conditionally independent given the nodes that connect them. Intuitively, this means (for example) that the relationship between “Daughter 1” and “Daughter 2” is entirely mediated by their shared relationship with “Mother 1”. Applying the DPI, the Markov assumption would imply

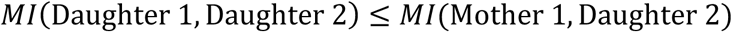

Thus, a violation of the above inequality, by contradicting the assumption of Markovity, would imply the existence of hidden properties. Due to the high-dimensional nature of scSeq data, as well as the continuity of early progenitor cell states, we found that calculating mutual information directly was impractical. Therefore, we took the alternative approach of applying machine learning algorithms to ask how well “Daughter 1” and “Mother 1” could respectively predict the state of “Daughter 2”. The intuition is that higher prediction accuracy would imply greater shared information about state.

### Data analysis for assessment of hidden properties

Following the theoretical discussion above, the existence of hidden properties could be inferred from the following inequality:

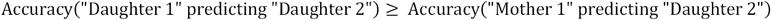

Since most machine learning algorithms perform discrete classification based on continuous, high-dimensional input data, we specifically assessed how well the gene expression states of “Daughter 1” and “Mother 1” could respectively predict the discrete lineage identity of “Daughter 2”. In our data, “Mother 1” and “Mother 2” would refer to clonally related cells from day 2, and “Daughter 1” and “Daughter 2” would refer to clonally related cells found in different wells on day 6 (the cells need to have been plated in separate wells on day 2 to ensure that their most recent common ancestor existed before day 2). Note that the terms “mother” and “daughter” are used for simplicity to refer to the early and late time points, but there is no requirement that mother cells divide.

We constructed training sets for each machine learning algorithm as follows. For each clone that appeared in two separate wells on day 6, we recorded the most common fate of the clone in one well, and the average transcriptome of the clone in the other well. Likewise, for each clone that appeared on day 6 and day 2, we recorded the most common fate of the day 6 cells and the average transcriptome of the day 2 cells. In each case, we then asked how well the most common fate could be predicted from the average transcriptome by applying a panel of machine learning algorithms (see below). We performed 100 splits into training and testing sets, and show the average accuracy of each algorithm in Fig. 5c, where accuracy is defined as the fraction of correct guesses for most common fate.

For machine learning algorithms, we applied the following functions from the python 2.7 module sklearn (version 0.19.2). The parameters used for each function are also shown below. All parameters were default except for the “hidden_layer_sizes” parameter of the MLPClassifier, where we used the non-default value of (50,20).

`sklearn.ensemble.RandomForestClassifier` (*n_estimators=‘warn’*, *criterion=‘gini’*, *max_depth=No ne*, *min_samples_split=2*, *min_samples_leaf=1*, *min_weight_fraction_leaf=0*.*0*, *max_features=‘auto’*, *max _leaf_nodes=None*, *min_impurity_decrease=0*.*0*, *min_impurity_split=None*, *bootstrap=True*, *oob_score=F alse*, *n_jobs=None*, *random_state=None*, *verbose=0*, *warm_start=False*, *class_weight=None*)

`sklearn.neural_network.MLPClassifier` *(hidden_layer_sizes=(50*, *20)*, *activation=‘relu’*, *solver=‘adam’*, *alpha=0*.*0001*, *batch_size=‘auto’*, *learning_rate=‘constant’*, *learning_rate_init=0*.*001*, *power_t=0*.*5*, *max_iter=200*, *shuffle=True*, *random_state=None*, *tol=0*.*0001*, *verbose=False*, *warm_start=False*, *momentum=0*.*9*, *nesterovs_momentum=True*, *early_stopping=False*, *validation_fraction=0*.*1*, *beta_1=0*.*9*, *beta_2=0*.*999*, *epsilon=1e-08)*

`sklearn.neighbors.KNeighborsClassifier` (n_neighbors=5, weights=‘uniform’, algorithm=‘auto’, leaf_size=30, p=2, metric=‘minkowski’, metric_params=None, n_jobs=1, **kwargs)

`sklearn.naive_bayes.GaussianNB` *(priors=None)*

## Application of fate prediction algorithms

### Classification of neutrophil/monocyte trajectory cells

Putative neutrophil/monocyte trajectory cells were identified using a stepwise process of graph smoothing and thresholding. Briefly, indicator variables representing each mature lineage as well as MPPs (see the previous methods section “Graph connectivity score” for a definition of indicator variables) were smoothed (N=250 smoothing iterations) and summed together to form an aggregate score. In the summation, smoothed scores for monocytes, neutrophils and MPPs were given a positive coefficient, and the other lineages got a negative coefficient. Formally, let *S*_*ij*_ denote the smoothed score for the *j*-th lineage in the *i*-th cell, and let *a*_*i*_ be the coefficient for the *j*-th lineage. Then the aggregate score *Z* was computed for each cell as 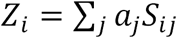, with the following values for *a*_*i*_: Neu=0.1, Mo=0.1, MPP=1, Meg = −2, All other lineages = −1.

The aggregate score (*Z*) was then thresholded to generate an approximate indicator *I*_*i*_ of the neutrophil/monocyte trajectory. The threshold was chosen manually to maximize the number of Neu/Mo trajectory cells while excluding cells that appeared to be differentiating toward other lineages. The final value resulted in 40,000 cells passing threshold. Because the indicator was noisy, another round of smoothing was performed on the indicator (N=50) and then the smoothened indicator was re-thresholded (threshold = 0.6) to generate a final annotation of neutrophil/monocyte trajectory cells.

### Population Balance Analysis

We performed PBA as described in (Weinreb, Wolock et al. 2018), using the python pipeline available at (https://github.com/AllonKleinLab/PBA). PBA was performed on a random 20,000-cell subset of the full dataset to limit the computational run-time. To extend the predicted fate probabilities to all cells, a 50-nearest-neighbor graph was constructed on the full dataset, and each cell inherited the mean value assigned to the nearest neighbors that were part of the 20,000-cell subset.

### Waddington-OT

We performed Wadding-OT analysis (Schiebinger, Shu et al. 2017) on the neutrophil/monocyte trajectory cells (Fig 5e) and on the whole dataset (Supp. Fig 7c) using python code available online (https://github.com/broadinstitute/wot). We did the analysis on a subset of cells (20,000 for neu/mo, 40,000 for all lineages) and then extended fate probabilities to the full dataset by averaging over 50 nearest neighbors (see above). The input to Wadding-OT is a set of time-point labeled transcriptomes and a proliferation score for each cell, and the output is a transition map between each pair of consecutive time points. We calculated proliferation scores in two steps. First, we used our state-fate barcoding data to calculate the clonal expansion downstream of each barcoded cell. Since these labels were very noisy, we then applied graph smoothing (N=15 rounds). Using the resulting proliferation scores, we ran Wadding-OT with parameters *λ*_1_ = 50 and *ϵ* ranging from 0.0001 to 50, where *λ*_1_ and *ϵ* are regularization parameters that control the fidelity of proliferation score constraints and the entropy of optimal transport map respectively. The output was a pair of transition maps *T*_2,4_ and *T*_4,6_ from day 2 to day 4 and day 4 to day 6 respectively, where 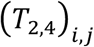 denotes the total density transported from cell *i* to cell *j*.

Terminal fate probabilities for each day 2 cell were calculated by (1) Row normalizing the *T*_2,4_ and *T*_4,6_ to generate transition probabilities; (2) Composing the normalized transition maps; (3) Collapsing the target cells of the transition map by lineage; (4) Linearly rescaling each lineage probability to ensure a uniform distribution over lineages among MPPs, to be consistent with PBA. Formally, these steps involved the following operations:

**Step 1: Row normalizing the** *T*_2,4_ **and** *T*_4,6_

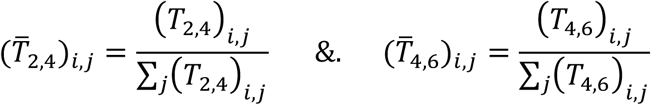

**Step 2: Composing the normalized transition maps**

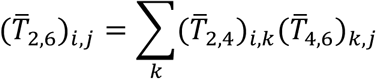

**Step 3: Collapsing the target cells of the transition map by lineage**

Let *X*_*ij*_ be an indicator variable where *X*_*ij*_ = 1 if cell *i* is in lineage *j*, otherwise *X*_*ij*_ = 0. We calculate *P*_*ij*_, the probability that cell *i* (from day 2) gives rise to lineage *j*, as follows:

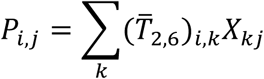

**Step 4: Linearly rescaling each lineage probability to ensure a uniform distribution over lineages among MPPs**

Let *X*_*i*_ be an indicator variable where *X*_*i*_ = 1 if cell *i* is an MPP, otherwise *X*_*i*_ = 0, and let *Q_j_* 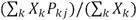 be the average predicted probability for an MPP to give rise to lineage *j*. Starting with lineage probabilities *P*_*ij*_, we calculate rescaled probabilities 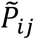 as follows:

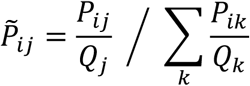

### FateID

We performed FateID analysis (Herman, Sagar et al. 2018) using the R package (https://github.com/dgrun/FateID). We did the analysis on a subset of 20,000 cells and then extended to the full dataset. The top 1000 most highly variable genes were used. As with PBA and Wadding-OT, lineage probabilities were linearly rescaled to ensure a uniform distribution of fates for MPPs (see step 4 above).

## Pseudotime analysis

### Classification of neutrophil trajectory cells

To classify cells as belonging to the neutrophil trajectory, we followed a similar procedure to that described above in section “Classification of neutrophil/monocyte trajectory cells”, but with the following weights. *a*_*i*_: Neu=0.1, MPP=0.1, Meg = −2, All other lineages = −1.

### Pseudotime ordering assignment

We assigned a pseudotime value to each cell in the neutrophil trajectory using graph smoothing. An initial indicator vector with 1 for MPP cells and 0 for all other cells was smoothed (N=300 rounds), generating a gradient of values that was highest near the MPPs and decreased along the trajectory. These values were quantile normalized to generate a final set of pseudotimes.

### Calculating the timing of differentiation for a typical cell

Let *t* represent real (calendar) time, and *F* represent pseudotime. To calculate the function *F*(*t*) from clonal barcoding data (Fig 5f), we began by calculating the average change in pseudotime over 2 days for cells starting at different points in the trajectory. The resulting function 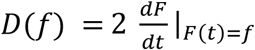, is proportional to the derivative of *F* with respect to *t*, written as a function of *F*. To transform D(f) into F(t), we can use the following general formula for taking the derivative of an inverse function:

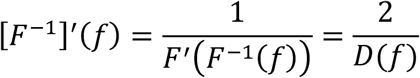

It follows that

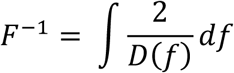

And therefore

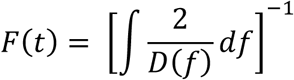

In practice, we calculated *D*(*f*) by first collecting all pairs of clonally related cells that appear two days apart and recording the change in pseudotime (*ΔT*) between them, as well as the pseudotime *f* of the starting cell. Note that, 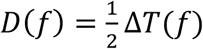. Since the *ΔT* values were noisy, we used curve fitting to estimate *D*(*f*). Curve fitting was performed in python using the function UnivariateSpline from package scipy.interpolate, with smoothing factor = 10^9^ and all other parameters default. We then numerically integrated *D*(*f*) to arrive at an estimate for *F*(*t*).

## Analysis of monocyte ontogeny with sleeping beauty transposon system

### Labeling of Sleeping Beauty Transposon Mice

The Sleeping Beauty Transposon mouse model was used as described previously (Sun et al. 2014, Rodriguez-Fraticelli et al. 2018). Briefly, we crossed mice carrying homozygous knock-in insertions of the DsRed2-Sleeping Beauty Transposon reporter in the *colla1* locus (Tn/Tn) and mice carrying homozygous knock-in insertions of the Hyperactive Sleeping Beauty Transposase in the *col1a1* locus (SB/SB) and knock-in insertion of the M2-rtTA tetracyclin-responsive transactivator in the *Rosa26* locus (M2/M2) to generate barcodable *col1a1*^*SB/TnRosa26M2rtTA/+*^ mice. Two 3 month-old mice were labeled by 2mg/ml Doxycycline-hyclate (Sigma-Aldrich) with 5mg/ml sucrose in drinking water for 1 week. Thereafter, Dox was removed and successful labelling (∼15% DsRed) was verified by retroorbital sinus peripheral blood collection (70 µl) after 1 week. Mice were euthanized 12 weeks after labeling. All animal procedures were approved by the Boston Children’s Hospital Institutional Animal Care and Use Committee.

### Bone Marrow and Spleen Cell Isolation

After euthanasia, whole BM (excluding the cranium) and spleen were dissected and processed separately. Cellular fractions were prepared in 2% fetal bovine serum in phosphate buffered saline, filtered with 70 µm filters, and then erythrocytes were removed with red blood cell lysis buffer. Lineage depletion was performed using Magnetic Assisted Cell Separation (Miltenyi Biotec) with anti-cKit magnetic beads. The cKit+ fraction was stained with CD117 APC (cKit, ACK2, Biolegend, 1:100), Ly6a PE/Cy7 (Sca-1, D7, eBiosciences, 1:100), CD34 FITC (RAM34, eBiosciences, 1:50), CD135 PE/Cy5 (Flt3, A2F10, eBiosciences, 1:50), CD115 BV605 (CSF-1R, AFS98, Biolegend, 1:100), CD48 APC/Cy7 (HM48-1, eBiosciences, 1:100), CD3e eFluor450 (145-2C11, eBiosciences, 1:100), CD19 eFluor450 (MB19-1, eBiosciences, 1:100), Ter119 eFluor450 (TER119, eBiosciences, 1:100), Gr1 eFluor450 (RB6-685, eBiosciences, 1:100) and CD11b eFluor450 (Mac1, M1/70, eBiosciences, 1:100). The fractions sorted were MPP (Lin^−^cKit^+^Sca1^+^CD48^+^), EryP (Lin^−^cKit^+^Sca1^−^CD34^−^FcgRII/III^−^Flt3^−^), GMP (Lin^−^cKit^+^Sca1^−^CD34^+^CD115^lo^FcgRII/III^+^Flt3^−^) and MDP (Lin^−^cKit^+^Sca1^−^ CD34^+^CD115^+^FcgRII/III^lo^Flt3^+^). The cKit-fraction was stained with CD115 BV605 (CSF-1R, AFS98, Biolegend, 1:50), Biolegend, 1:100), Ly6C APC (HK1.4, Biolegend, 1:200), Ly6G-AF700 (1A8, eBiosciences, 1:100), Ter119 eFluor450 (TER119, eBiosciences, 1:100), CD19 APC/Cy7 (1D3, eBiosciences, 1:100), Cd11c FITC (HC3, Biolegend, 1:100), CD74 BV711 (MHC-II, ln-1, Biolegend, 1:100) and CD43-PECy7 (S11, Biolegend, 1:100). Classic Monocytes were isolated as CD115^+^CD19^−^Ter119^−^Ly6C^hi^Ly6G^−^CD43^−^ cells from the bone marrow, Neutrophils were isolated as CD115^−^CD19^−^Ter119^−^Ly6C^lo^Ly6G^+^CD43^−^CD74^−^ cells from the bone marrow, and DCs were isolated as CD115^lo^CD19^−^Ter119^−^Ly6C^−^Ly6G^−^CD11c^hi^CD74^+^ cells from the spleen. For Tn tag content extraction and analysis, only DsRed2^+^ cells were sorted from each fraction. We FACS-sorted all the available cells from the whole BM or spleen extract using purity modes (∼95% purity) at ∼75–80% sorting efficiency.

### Transposon integration retrieval and analysis

Cells of interest were sorted into 1.7 ml tubes and concentrated into 5–10 µl of buffer by low speed centrifugation (700 g for 5 minutes). Transposon insertion site retrieval and analysis was performed with an adapted version of TARIS (Rodriguez-Fraticelli et al. 2018). Library indexing was used with Illumina library construction kit primers and sequencing was carried out on NextSeq (Illumina) at the Biopolymers Facility (Harvard Medical School). Tag identification and alignment was performed with a custom script as previously described (Sun et al. 2014). Briefly, we grep the Tn-containing reads from each fastq file, trim the adaptor and Tn sequences, and align the integration sites to the reference mouse genome (Ensembl mm9) using bowtie 1.2.

Samples with fewer than 10,000 mapping reads were discarded. Then, reads are normalized between samples (per million reads) and compared with at least one additional independently labelled mouse with libraries prepared in parallel and sequenced in the same NextSeq lane to account for contaminations. Tags present in the control mouse samples or only in one of the split samples were filtered out (contaminating reads). Next, read frequencies were log normalized and plotted using a heatmap (rows: unique barcodes, columns: sorted cell populations). Custom primers used were: Tn1-C primer (5′-CTT GTG TCA TGC ACA AAG TAG ATG TCC-3′), MAF-Tn1-1F primer (5′-ACA CTC TTT CCC TAC ACG ACG CTC TTC CGA TCT NNN NCG AGT TTT AAT GAC TCC AAC T-3′), and MAR-LCII (5’-GTGACTGGAGTTCAGACGTGTGCTCTTCCGATCTAGTGGCACAGCAGTTAGG-3’). All primers were ordered from IDT DNA technologies, at 100 nmole scale and HPLC-purified.

## SUPPLEMENT

**Supplementary Figure 1:**
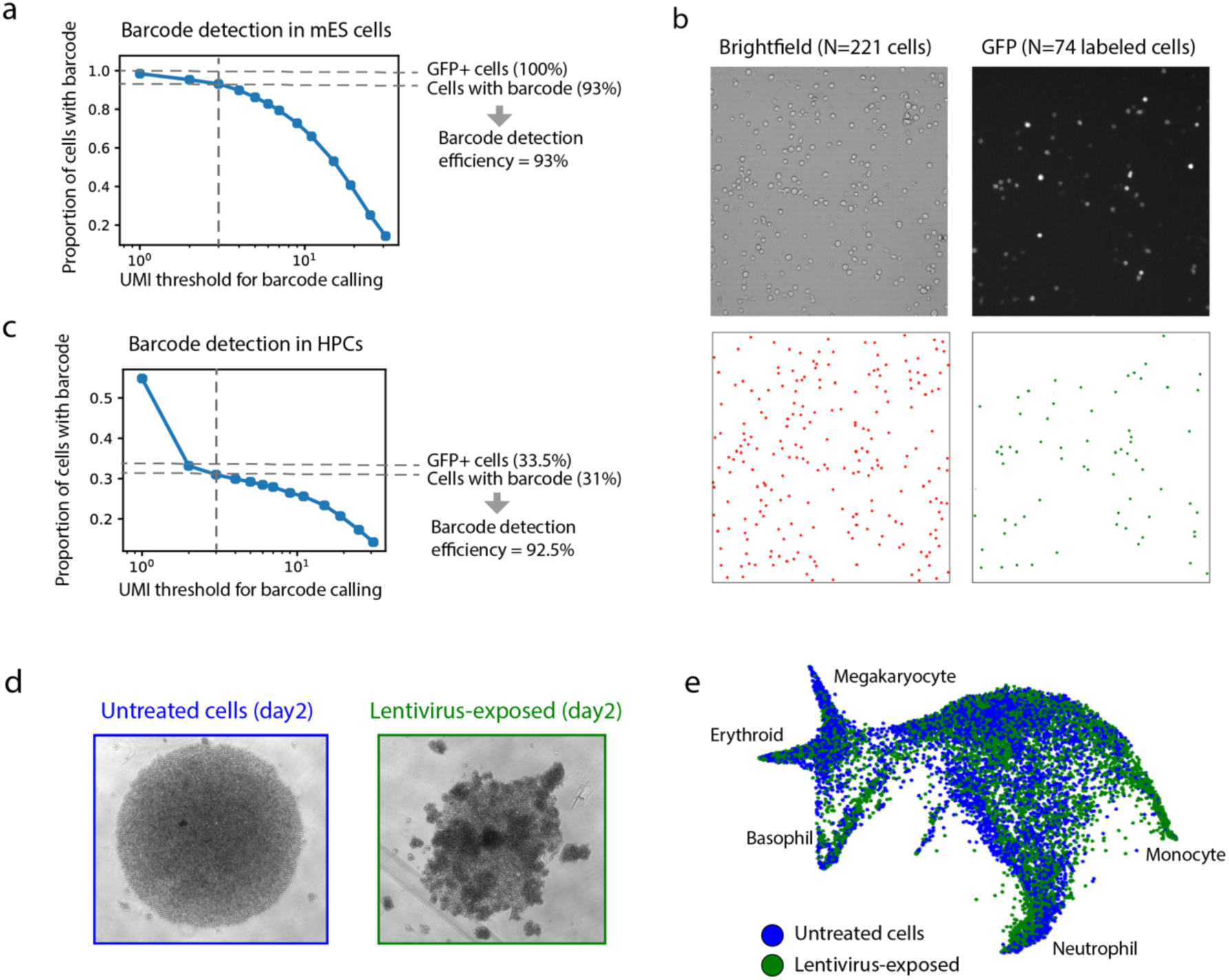
Lineage and RNA Recovery (LARRY) detection efficiency and effect on cells. (a) Proportion of cells with *> n* UMIs of a LARRY barcode as a function of *n* in GFP-sorted embryonic stem cells. Dashed line indicates filtering threshold to accept a LARRY barcode (minimum 3 UMIs). (b) Microscopy-based measurement of viral transduction efficiency in HPCs. Cell counts in brightfield and FITC images (top) are computationally scored (bottom) to measure the fraction of GFP+ cells (74/225 cells). (c) Proportion of cells with *> n* UMIs of a LARRY barcode as a function of *n* in HPCs from the same pool shown in panel (b). Dashed line threshold as in (a). (d) Brightfield images of virus-transduced HPCs (right) and naïve HPCS (left) in a round-bottom well, showing transient altered cell adhesion in transduced cells. (e) scRNA-Seq analysis to examine the effect of lentiviral treatment of HPCs, shown by a SPRING plot of treated and untreated HPCs [the same HPCs shown in panel(d)]. The two conditions show a similar distribution of gene expression states.

**Supplementary Figure 2:**
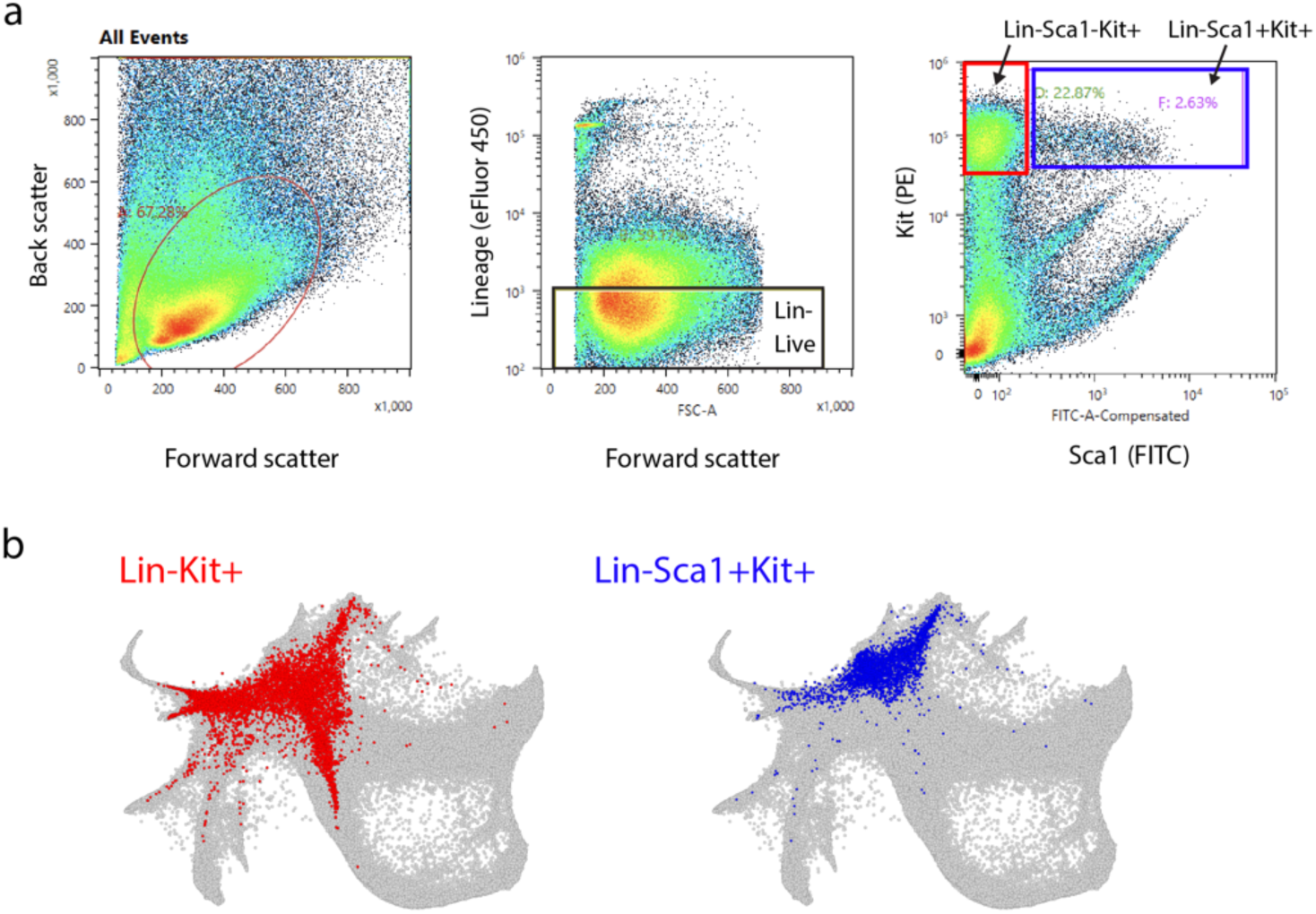
Isolation of progenitor cell populations. (a) FACS gates used to sort Lin-Kit+ (LK) and Lin-Kit+Sca1+ (LSK) cells. (b) LK and LSK projected onto the SPRING plot (from Fig. 1e) of cells from all time points of in vitro hematopoiesis, showing that LSK cells are less differentiated but are still a heterogeneous population.

**Supplementary Figure 3:**
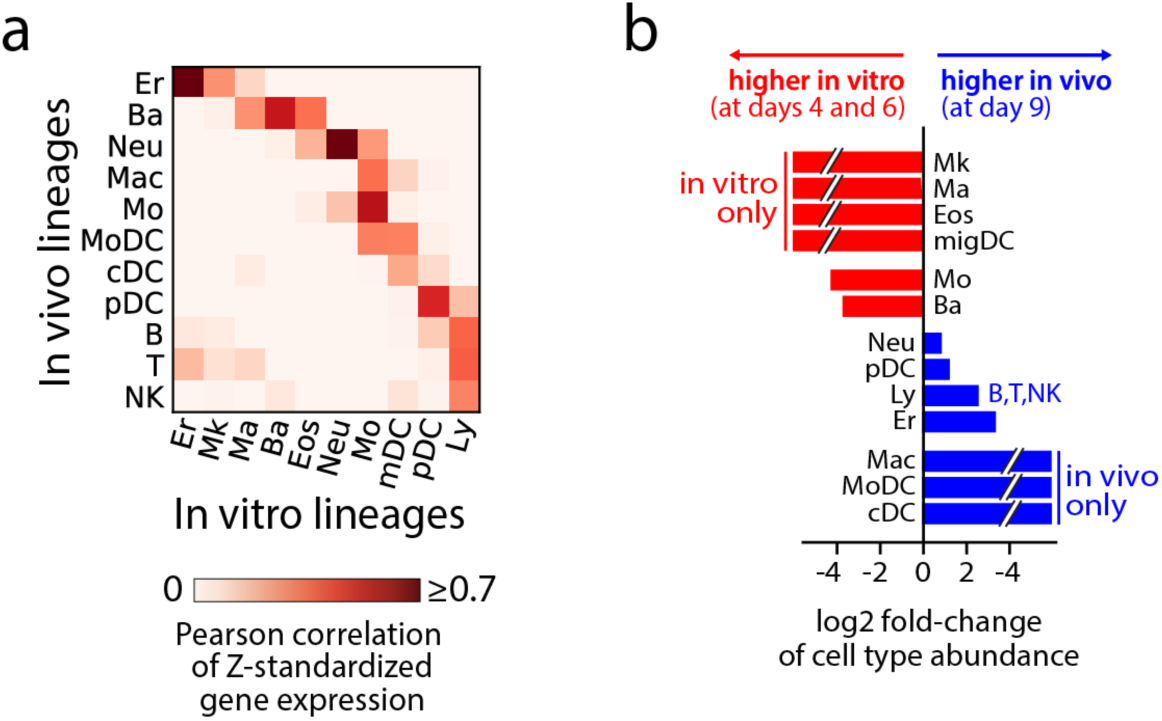
Cell states one-week post-transplantation. (a) Comparison of gene expression between mature cells in vitro and post-transplantation. Rows represent transplanted cell states and columns represent cell states in vitro. Color shows the Pearson correlation of Z-scores gene expression, restricted to highly variable genes. “Mature cells” are defined as cells within colored regions of the SPRING plot in Figs. 1e,f. These regions correspond to cells expressing high levels of genes associated with mature cell types (see methods for region delineation, and Table 2 for genes used). (b) Cell type abundances in vitro and after transplantation.

**Supplementary Figure 4:**
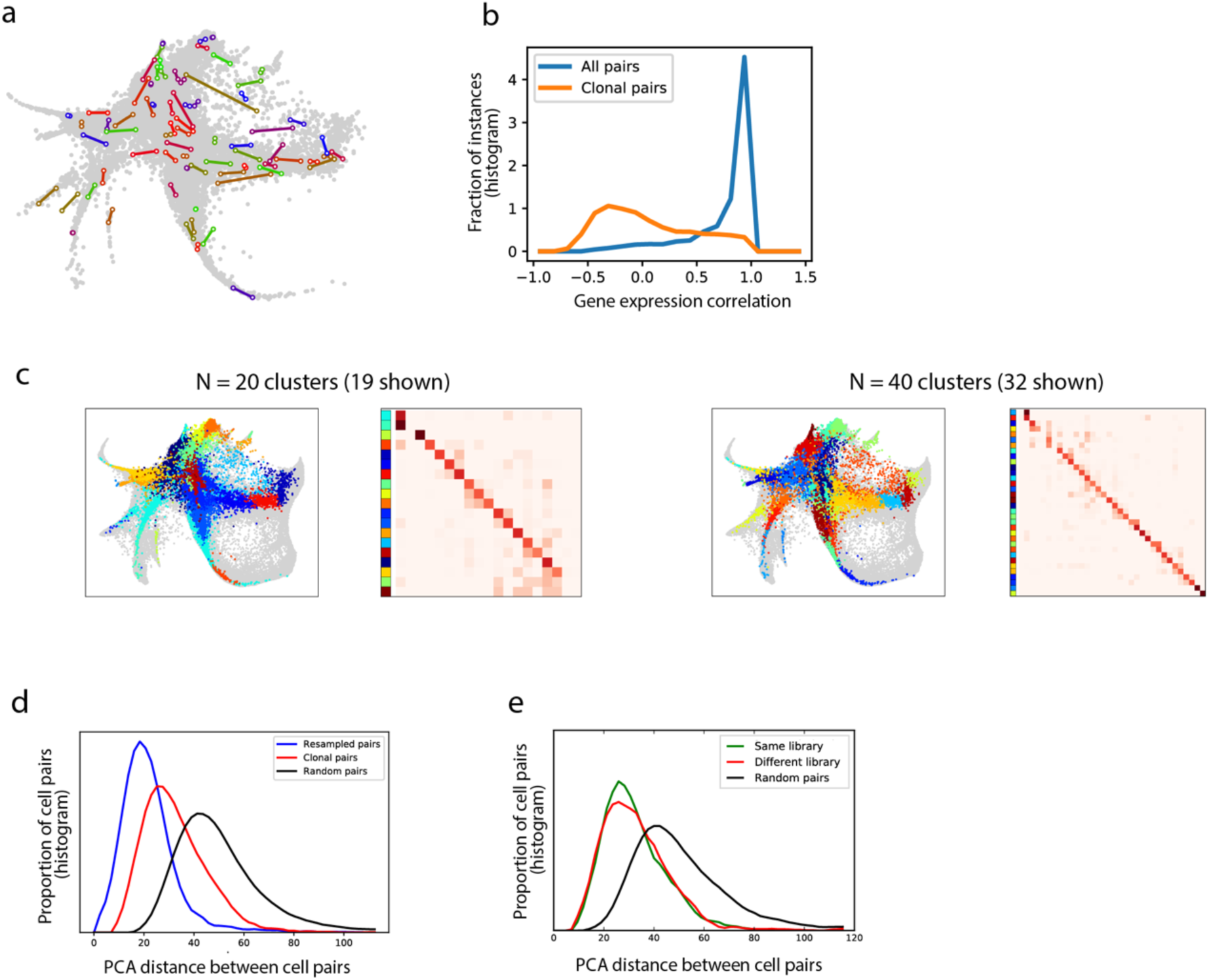
Analysis of sister cell differences on day 2. (a) SPRING plot visualization of a sample of sister cell pairs 2 days post-barcoding. Each colored line spans between two sister cells. The sister cell pairs were sampled so that they would be non-overlapping, but also have 2D distances representative of the full distribution. Grey points show the full data set across all time points. (b) Histograms showing correlation of gene expression for sister cells (blue line) versus random pairs (orange line). (c) Co-localization of sister cells in clusters shown for two different clustering granularities. In each case, the cluster labels for each cell are plotted on the left, and a heatmap showing the joint probability for a pair of sisters to appear in two different clusters is shown on the right. Out of N total clusters, only a subset is shown in the heatmap, since we excluded clusters with <0.1% of the cells. (d) Distribution of high dimensional distances between: random cell pairs (black); sister cells (red); and simulated cell pairs that differ only as a result of binomial sampling of transcripts (blue). (e) Distribution of high dimensional distances between: random cell pairs (black); sister cell pairs separated and subject to separate inDrop cell encapsulation runs (red); and sister cell pairs from the same library (green). The difference between the green and red curves indicates the degree to which sister cell similarity is driven by single-cell encapsulation artifacts such as gel doublets.

**Supplementary Figure 5:**
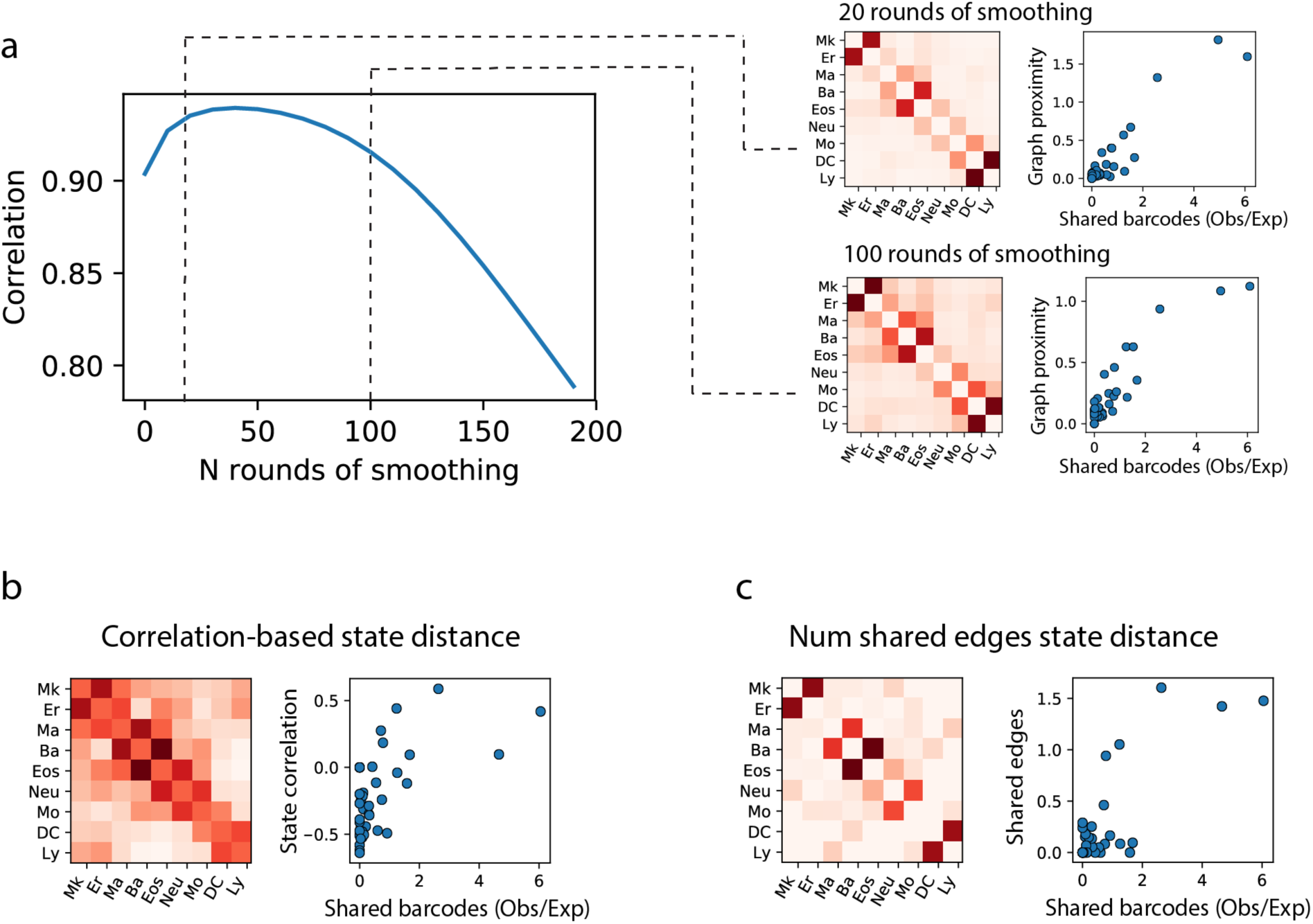
Robustness analysis for the comparison of state and fate distance in in vitro hematopoiesis. (a) Sensitivity analysis showing that correlation between fate distance and state distance remains stable across graph-smoothing iterations in the calculation of state distance. Correlation predictably drops at high iterations as all states become equally close. Blow-outs show a heat map of state distances (left) and their relationship with fate distances (right) for N=20 and N=100 smoothing rounds. (b) Heatmap of correlation-based state distances (left) and their relationship with fate distances (right). Note that this metric directly compares gene expression similarity, which is expected to be a less reliable indicator of developmental relationship than the graph connectivity via transitional states. (c) State distance based on number of shared edges between lineages in a nearest neighbor graph (left) and their relationship with fate distances (right). Note that this distance corresponds to N=1 iterations in panel (a).

**Supplementary Figure 6:**
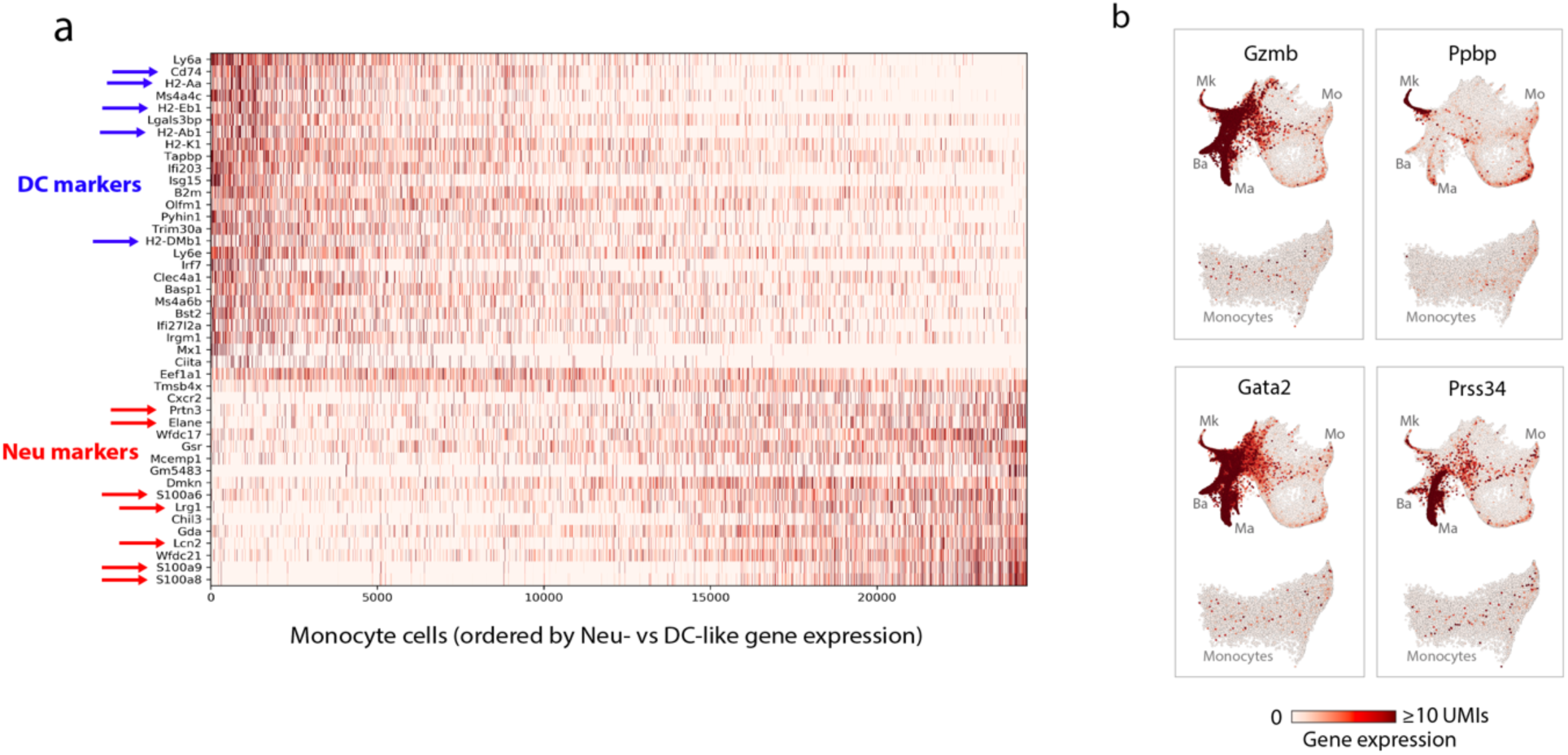
Heterogeneity in the gene expression and ontogeny of monocytes. (a) Heatmap showing expression of genes enriched among DC-like monocytes (top 20 rows) or among Neu-like monocytes (bottom 20 rows). Known marker genes for DCs or neutrophils are indicated by arrows. References are provided in Table 2. (b) Expression of basophil, mast cell and megakaryocyte marker genes in the whole dataset (top plot in each panel) or just monocytes (bottom plot in each panel) shows that only neutrophil and DC markers are enriched among monocytes. Since basophil, mast cells and megakaryocytes combined are 70% as abundant as neutrophils, the neutrophil and DC marker gene expression in monocytes is unlikely to be caused by cell doublets, which would affect all lineages equally.

**Supplementary Figure 7:**
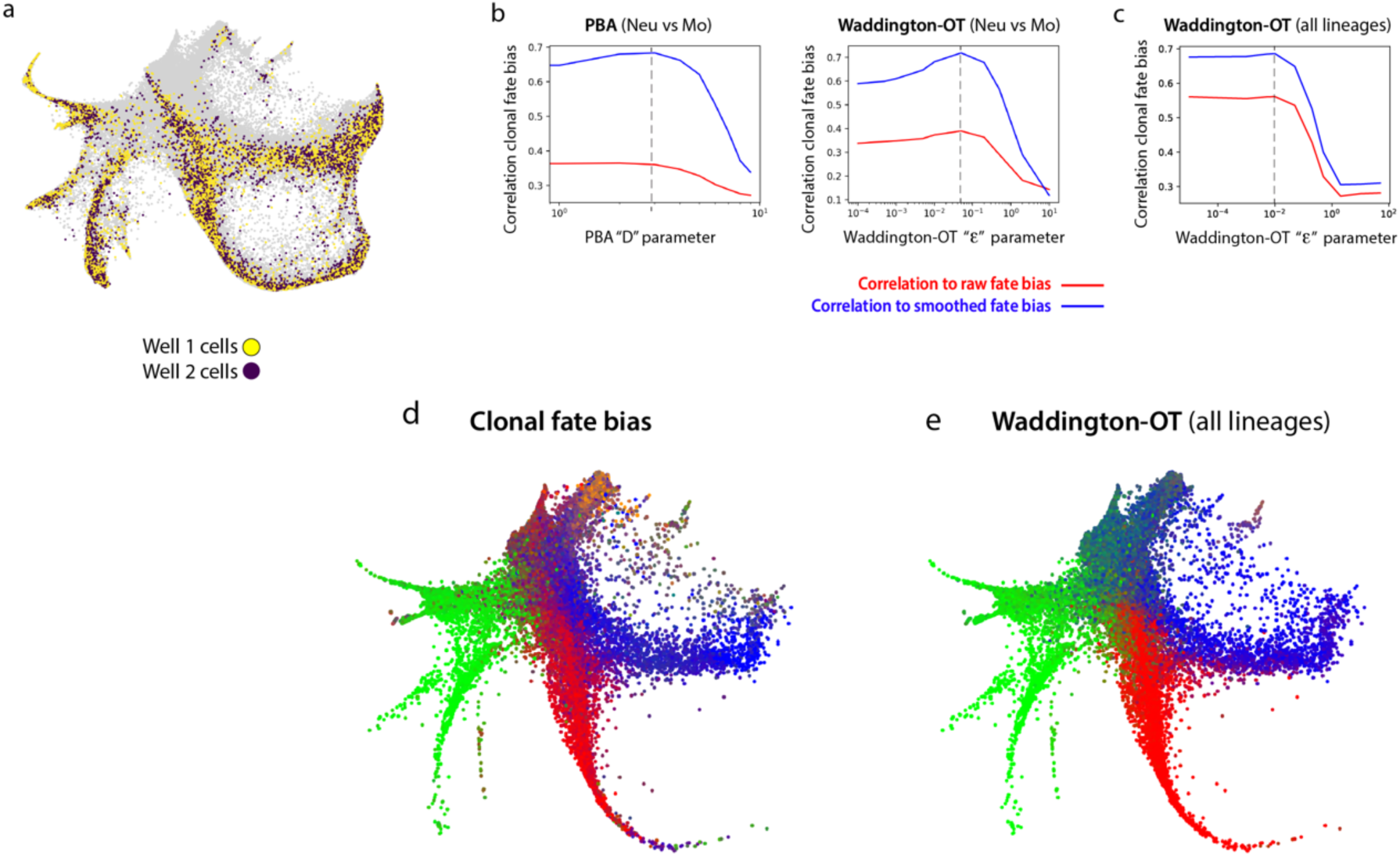
Prediction of fate probability from transcription alone. **(a)** Locations of day 6 cells cultured into two separate wells, used for analysis of hidden properties influencing cell fate. (b) Parameter scans of computational fate prediction algorithms to identify their optimal theoretical performance as measured by correlation between predicted neutrophil vs monocyte fate probability and empirical clonal fate probability. The PBA diffusion (*D*) parameter and WADDINGTON-OT entropy (*ϵ*) parameter are scanned. Dashed lines indicate optimal values used for comparison. (c) Same as panel (b), right, but now for all lineages. (d) Empirical clonal fate bias represented by a multi-color mixture on a SPRING plot of cells from day 2. Pure colors indicate graph regions where all clones have the same fate outcome. Gata1/2+ fates (erythroid, megakaryocyte, basophil, eosinophil and mast cells) are pooled together for this visualization. (e) Waddington-OT predicted fate bias shown on a day 2 SPRING plot.

**Supplementary Figure 8:**
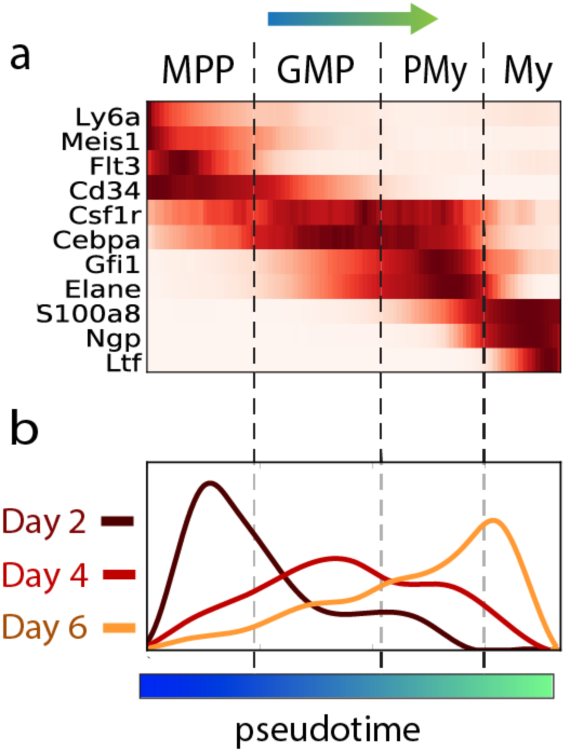
Supporting data for pseudotime progression analysis. (a) Marker genes used to establish boundaries between stages [MPP=multipotent progenitor; GMP=granulocyte monocyte progenitor; PMy=promyelocyte, My=myelocyte]. (b) Distribution of pseudotimes for each time point. Cells within a time point are asynchronous, but accumulate toward the end of the trajectory over time.

**Supplementary Figure 9:**
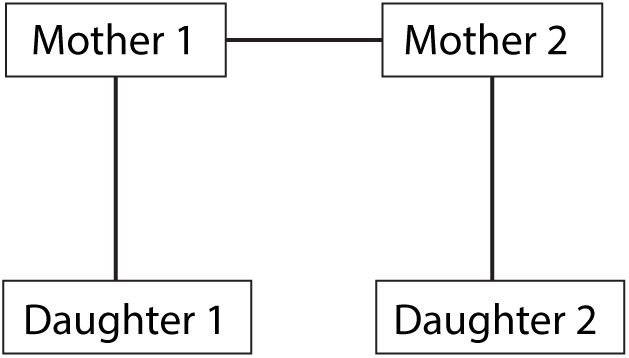
Graphical model showing dependencies between clonally related cells at different time points.

